# Knowledge gap and conditional receptivity: why tree and forest professionals underutilize citizen science for quarantine pest detection in France

**DOI:** 10.64898/2026.07.30.740952

**Authors:** Bastien Castagneyrol, Souleymane Bah, Baptiste Bedessem, Maarten de Groot, Amal Ennabih

## Abstract

Exotic pests and pathogens pose a major threat to forest ecosystems. Early detection of newly introduced organisms is critical for implementing effective eradication measures before they become established. Citizen science platforms have emerged as promising sources of biodiversity data for detecting invasive pests and pathogens, yet their integration into formal biosecurity surveillance remains limited. Using France as a case study, we surveyed 101 professionals involved in forest and urban tree management and pest surveillance to assess their knowledge of regulated tree pests and pathogens and their attitudes towards online citizen science platforms for biosecurity applications. We focused on ten focal species representing different regulatory statuses under EU legislation. We assessed the knowledge professionals have of these species and their regulatory status, and explored sources of variation in their opinions regarding the use of citizen science platforms as a tool for post-border biosecurity. Experts involved in mandatory surveillance of regulated organisms (SORE experts) demonstrated consistent knowledge of quarantine pests, whereas other professionals’ knowledge varied by species, with greater familiarity for non-quarantine, widely distributed pests. Half of respondents used citizen science platforms, predominantly to consult species distribution rather than to contribute observations. Professionals’ receptivity to citizen science increased significantly when species were perceived as easy to identify, but they expressed more confidence in citizen science for monitoring established pests than for the early detection of quarantine species. The survey reveals that while citizen science platforms are known and valued as information sources, they remain underutilized as mechanisms for sharing field observations, even among surveillance professionals.

## Introduction

Forests are at risk. Whereas planted, natural, and urban forests are at the cornerstone of attenuation and adaptation to climate change (Bonan 2008; Escobedo et al. 2019; Savo et al. 2025; Reek et al. 2026), the sustainability of the functions and services they provide to humanity is increasingly threatened. Climate change fuels the intensity and frequency of major disturbances responsible for forest loss—including drought, fires, and storms—but also chronically increases their vulnerability to pests and diseases (Seidl et al. 2017; Pureswaran et al. 2018; Jactel et al. 2019; Hlásny et al. 2025). International trade further amplifies biotic risk by moving potentially harmful species beyond their native range, thus creating opportunities for novel interactions between trees, pests, and pathogens (Liebhold et al. 2023). The biological causes and consequences of the biotic and abiotic risks to which forests are exposed are increasingly well documented, as are the economic drivers of biological introductions (Ramsfield et al. 2016; Nahrung et al. 2023). However, the social dimension of forest vulnerability has been largely overlooked to date (Marzano et al. 2016; Raum et al. 2024).

Biosecurity surveillance is an essential activity designed to discover non-native species before their introduction, or at the earliest stage of biological invasion (Nahrung et al. 2023). When the introduction of an exotic pest or pathogen is detected soon enough after introduction (i.e. post-border surveillance) or even before its introduction to a new area (i.e. pre-border surveillance), the probability of successfully implementing eradication or containment measures increase (Tobin et al. 2014; Branco et al. 2023), thus calling for the development of effective early warning schemes. The European Commission has established a list of quarantine species whose active surveillance by national authorities is mandatory across the European Union, and whose detection triggers a series of reporting and eradication procedures aimed at eradicating the pest. Despite substantial regulatory, human, and financial resources deployed to this end, a recent study revealed that eradication attempts for invasive pests and pathogens concerned only a minority of biological introductions (Branco et al. 2023). Active surveillance and regulatory tools are therefore insufficient to tackle the risk posed by non-native pests and pathogens (Klapwijk et al. 2016), particularly if the number of observers or their knowledge of the regulations is limited.

There is a continuum of vulnerability of trees across urban, planted, and natural forests (Epanchin-Niell and Pi 2024; Augustinus et al. 2024). Introductions of pests and pathogens often begin in close proximity to large cities, where their establishment is facilitated by the high diversity of trees, including non-native trees likely to serve as bridgeheads for biological invasions (Dodds and Orwig 2011; Paap et al. 2017; Branco et al. 2019). Whereas pests and pathogens can move freely across human habitat boundaries, biosecurity is, in practice, often siloed, with agencies in charge of plant health surveillance and protection in cities being different from those in charge of forest health surveillance (Tomlinson et al. 2015; Raum et al. 2025). In France, for instance, the surveillance of regulated and emerging pests (SORE, *Surveillance des Organismes Réglementés et Emergents*) is a centralised, top-down implementation of EU Regulation 2016/2031 built on a multi-level, multi-actor configuration and legally binding, standardised data-collection circuits. The SORE is operated by plant protection officers belonging or closely related to the Ministry of Agriculture. SORE experts are commissioned every year to inspect plants in specific habitats (forests, agricultural lands, urban green spaces, infrastructures and roadsides), searching for the presence of a pre-defined list of pests and pathogens, including those classified as quarantine organisms under EU Regulations (EU) 2019/2072 and (EU) 2019/1702. However, agents and agencies operating in forest have little connections with those operating in cities our the agricultural sector. It therefore appears necessary to develop biosecurity approaches that transcend these geographical and administrative boundaries in order to better reflect biological and ecological realities.

Citizen science can reinforce biosecurity surveillance by increasing the number of vigilant eyes in the field (Baker et al. 2019; Gupta et al. 2022; Hulbert et al. 2023; de Groot et al. 2023). Citizen science is an umbrella term describing any form of contribution by people who are not professional scientists to scientific research. It includes the general public as individuals or organized groups as well as professionals from specific sectors who may have an interest in a particular research area (Haklay et al. 2021). In terms of biosecurity, participation in citizen science contributes to raise public awareness of invasive alien species and the problem they poses while advancing scientific knowledge (Pocock et al. 2024). It ranges from structured and standardized surveillance schemes to semi-structured bioblitzes, or incidental contributions through the sharing of pictures of insects on online naturalist platforms (Meentemeyer et al. 2015; Brown et al. 2020; Price-Jones et al. 2022; Meeus et al. 2023; Roe et al. 2024). Although opportunistic and unstructured, these latter forms of public contribution to the surveillance of invasive alien species have proven to be powerful tools for early detection, particularly for large and conspicuous species (Pocock et al. 2024). In several instances, the presence of an invasive pest in a country was first reported on online naturalist platforms before its presence was known to the national authorities (Epanchin-Niell et al. 2021; Roe et al. 2024; González-Moreno et al. 2025).

Essential to biosecurity are feedback loops and efficient information flows between people making field observations and those in charge of implementing pest and pathogen regulation schemes (Ticehurst and Kruger 2023; Hutt-Taylor et al. 2025). However, persistent doubts remain about the accuracy and value of data collected by observers whose expertise is unknown (Austen et al. 2016; Collins et al. 2022). Citizen science data indeed suffer from several well-known biases (Pocock et al. 2024): large and conspicuous species are often overrepresented compared with smaller or less conspicuous species (Caley et al. 2020), and observations tend to be concentrated where observers are —typically near urban areas— rather than where species occur (de Groot et al. 2022). These well-documented biases may explain experts’ reluctance to use data generated by laypeople. On the contrary, it may seem intuitive that any professional with daily and direct contact with plants through their activity (whether, for instance, arborists, forest or greenspace managers) would be the most concerned by tree health issues and the most likely to be the first observer of an introduced pest or pathogen. These professionals may thus constitute a particularly relevant target for biosecurity-oriented citizen science programmes (Thomas et al. 2017).

Despite the potential value of citizen science for health forest surveillance, the extent to which it is actually recognized and used by forest professionals is poorly documented. Using France as a case study, we designed a survey aiming to assess the knowledge and use of citizen science platforms among professionals working with trees (including urban trees) and forests, whether managers or plant protection officers involved in the surveillance of EU (henceforth ‘*SORE experts*’, or, in short, *‘experts’*). We hypothesized that SORE experts would have a better knowledge of regulated pests than other professionals, regardless of their sector of activity, because the work of SORE experts is to survey for these species, while other professionals focus on species that are harming city trees, regardless of their status. We further documented the level of knowledge they have of citizen science platforms and whether they use them, and if so for which purpose. We anticipated that the extent to which professionals value citizen science as a potential source of relevant information would depend on the conspicuousness of pests or disease symptoms. Specifically, we predicted that professionals would value citizen science data referring to large, easily identifiable species, and be more reluctant to consider citizen science as a valid tool for biosecurity for species whose identification requires greater expertise. By doing so, our study documents the social component of forest vulnerability to non-native pests and pathogens.

## Materials and methods

### Study site and model species

The study was conducted in France, Western Europe. The country covers 633 000 kmZ with a population approaching 70 millions inhabitants (INSEE 2026), 80% of each living in cities. Forests cover 25 millions hectares, i.e., 33% of the country (excluding ultramarine territories). Three quarters of it are privately owned. 39 Mm3 are harvested every year, representing a commercial value of 2800 M€ (Chuine et al. 2023), with 8% of trees being in sanitary state (IGN 2025).

We established a list of 10 focal tree pests and pathogens with different regulation statuses at the EU and national (France) levels **(Table 1)**. Specifically, we considered six quarantine species listed in Annex II of EU Regulation 2016/2031, of which two were not present in Europe at the time the survey was administered (*i.e.*, Annex IIA), and four were known to occur or have occurred within EU boundaries (*i.e.*, Annex IIB). We also considered four species that are regulated at the EU level but are not classified as quarantine species (*i.e.*, species in Annex III). The three insect species listed in Annex III are widely distributed in France and are frequently spotlighted in the mainstream media (Castagneyrol et al. 2026), either because of the major damage they cause to forests or because they can be harmful to humans. Specifically, the European spruce bark beetle is a major pest responsible for considerable damage to spruce forests. Its impact has intensified over the last decades as a consequence of the combined effects of forest mismanagement and climate change (Hlásny et al. 2021; Hallas et al. 2026). The pine and oak processionary moths are also important defoliators of pines and deciduous oaks, respectively, but are mostly known for their noxious urticating hairs (Battisti et al. 2015). We regarded these species as ‘control’ species that we expected to be well known to respondents. Also listed in Annex III, chestnut blight is a disease causing major damage to chestnut orchards and forests; it is widely distributed across the national territory, where it is not considered a quarantine species. Important to this study is the fact that, in November 2025, the first occurrence of the pine wood nematode was confirmed in southwestern France, attracting significant media attention.

**Table 1.** List of tree pests (i) and pathogens (p) included in the questionnaire. Respondents were presented scientific names and common names in French. Species listed in Annexes IIA and IIB are EU quarantine species, currently absent from the EU territory (IIA), or known to occur within the EU (IIB). Species listed in Annex III are regulated organisms that are not quarantine species at the EU level but are quarantine species in specific areas (though not in France). Note that as this article was being written (July 2026), the presence of the emerald ash borer (so far listed in Annex IIA) was confirmed in Hungary.

| French | Scientific | English | EU regulation status | Status in France* |
| --- | --- | --- | --- | --- |
| Nématode du pin | *Bursaphelenchus xylophilus* (Nematoda) | Pine wood nematod (p) | Annex II B | First detection in November 2025 in SW France |
| Agrile du frêne | *Agrilus planipennis* (Coleoptera) | Emerald ash borer (i) | Annex II A | Absent |
| Chancre résineux du pin | *Fusarium circinatum* (Ascomycota) | Pitch canker (p) | Annex II B | Absent, eradicated |
| Chancre du châtaignier | *Cryphonectria parasitica* (Ascomycota) | Chestnut blight (p) | Annex III | Present, restricted distribution |
| Processionnaire du chêne | *Thaumetopoea processionea* (Lepidoptera) | Oak processionary moth (i) | Annex III | Present, restricted distribution |
| Scolyte de l'épicéa | *Ips typographus* (Coleoptera) | European spruce bark beetle (i) | Annex III | Present, restricted distribution |
| Agrile du bouleau | *Agrilus anxius* (Coleoptera) | Bronze birch borer (i) | Annex II A | Absent |
| Processionnaire du pin | *Thaumetopoea pityocampa* (Lepidoptera) | Pine processionary moth (i) | Annex III | Present, expanding northward |
| Chancre coloré du platane | *Ceratocystis platani* (Ascomycota) | Canker stain of plane (p) | Annex II B | Present, restricted distribution |
| Capricorne asiatique | *Anoplophora glabripennis* (Coleoptera) | Asian longhorn beetle (i) | Annex II B | Present, restricted distribution |

We implemented an online survey between May July 2026. The survey was inspired by Raum et al. (2024) and was developed with the support of experts in forestry and urban green space management. It was intended to reach plant protection officers whose work involved trees, as well as various groups of professionals working with trees in forest, urban, and peri-urban environments. These categories included arborists and tree care takers, landscape planners and urban planners, park and green space managers and technicians, managers of public and private forests, and forest technicians and consultants. Respondents could identify themselves as belonging to more than one category. The first question asked respondents to confirm that they had or had had a professional activity related to trees.

The survey was administered using Limesurvey version 3. We used a targeted snowball sampling approach (Goodman 1961) to distribute the survey, sending formal invitations accompanied by brief background information to national contacts within key organisations **(Table 5)**. The invitation to participate in the survey was further disseminated through personal communications by colleagues from our research department who maintain well-established professional relationships with a wide range of stakeholders.

### Questionnaire design

The questionnaire (Appendix A, Supplementary Material) was divided into four sections and 16 questions (Q). The first section assessed respondents’ knowledge of the focal tree pests and pathogens (Q1, organized on a four-degree Likert scale), of their regulation status (Q2, yes/no question), and their degree of concern (Q3, four-degree Likert scale). The second section explored the use respondents might have of online citizen science platforms. We first asked whether respondents made any use of a pre-set list of five generalist and professionally oriented citizen science platforms (Q4): the generalist platforms *iNaturalist*, and *Observatoire des Espèces Exotiques Envahissantes*, the forest oriented platform *Silvalert*, and *Vigil’Encre* and *VigiCultures*, two platforms designed to survey the presence and spread of plant pests and diseases. Then, for each citizen science platform, we asked whether it was used in a personal or professional context (Q5) and, for professional use only, we asked why (Q6), how (Q7), what for (Q8) and how much (Q9) it was used.

The third section of the questionnaire aimed at understanding respondents’ attitudes regarding the potential use of online citizen science platforms for biosecurity. We first asked respondents’ opinion on the difficulty of identifying the focal tree pests and diseases (Q10), on the probability that users of citizen science platforms would share observations of these species (Q11) and on the probability that they —respondents— would consult citizen science platforms to obtain information regarding the distribution of each pest and pathogen (Q12). Q10-12 were on four-degree Likert scales.

The last section explored two scenarios designed to put respondents in situations in which they would use citizen science platforms in a professional context. The first scenario mentioned the Asian Longhorn Beetle *Anoplophora glabripennis*, a quarantine species that has occasionally be intercepted in France but has always been successfully eradicated so far, and the oak lace bug *Corythucha arcuata* a non regulated but invasive pest whose range has rapidly expanded seen its first detection in France in 2017. Scenarios read as follows: *“While preparing for an inspection mission, you consult a digital biodiversity oriented citizen science platform out of curiosity to learn about the species already present or reported in your area. While browsing the observations available on the platform, you come across a photograph showing* Anoplophora glabripennis/Corythucha arcuata, *although you have never observed this organism in this area before”*. Questions Q13-Q16 were on four-degrees Likert scales and asked respondents’ opinion on the possibility to obtain reliable abundance (Q13), absence (Q14), presence (Q16) and spatial distribution (Q16) of these two species in France. Eventually, we asked respondents their opinion regarding the potential of citizen science platforms as a relevant tool for tree-related biosecurity in France.

### Statistical analyses

We analysed data in R version 4.4.1 (Team 2024) using functions provided by the library `tidyverse` (Wickham et al. 2019) to summarized qualitative variables and visualized the data. The sample size was not sufficient to reliably analyse the detailed answers of respondents regarding their use of each citizen science platform (Q4-Q9). We regrouped their responses and eventually analysed respondents’ attitudes regarding citizen science platforms as a whole.

We analysed responses to Q1–Q3 with cumulative link mixed models (CLMMs), the ordinal analogues of ANOVA with a random effect. The response variable was the level of knowledge respondents had of each pest and pathogen, treated as an ordinal variable. In the first set of models, the explanatory variables were species identity and respondent status with respect to participation in the surveillance of regulated and emerging pests (expert *vs.* non-expert) as well as the two-way interaction. In the second set, we replaced species identity with a coarser functional group description (insect vs. pathogen) and with regulatory status under EU legislation (quarantine vs. non-quarantine). Respondent ID was included as a random effect to account for the possible non-independence of multiple responses from the same respondent. We used the same approach to test whether the characteristics of the pests and pathogens and respondent status influenced the probability of correctly identifying whether each organism is regulated. To this end, we treated each answer as an ordered response (incorrect < “don’t know”/missing < correct), considering a missing answer and “I don’t know” as equivalent because respondents went on to answer the subsequent questions. In all cases, we fitted CLMMs with the `clmm` function from the `ordinal` package (Christensen 2026), estimating parameters by adaptive Gauss–Hermite quadrature, and assessed the significance of each predictor with likelihood-ratio tests comparing nested models. We tested contrasts among factor levels with function emmeans from package emmeans (Lenth et al. 2026). In case there was a significant interaction between respondent status and pest features, we tested pairwise differences between pest categories within each level of respondent status.

To correlate respondents’ opinion on the difficulty in identifying pests and diseases with their beliefs about the likelihood of using citizen-science platforms to investigate the presence of pests and diseases in France and the likelihood that citizen scientists would report such pests and diseases on a platform (Q10-Q12), we transformed qualitative answers into weighted quantitative indices. Specifically, we assumed that the qualitative responses (very unlikely, unlikely, likely, very likely; very difficult, difficult, easy, very easy) reflected underlying continuous latent traits (levels of agreement) partitioned into four categories. We first recoded these answers on a scale from 1 to 4. Then, for each pest or disease, we multiplied each recoded score by the number of respondents selecting that level, summed the products across all levels, and finally divided the total by the overall number of responses. We finally regressed the scores obtained for the probability of reporting pest and disease observations on citizen-science platforms or the probability of consulting such a platform on the scores reflecting the difficulty of identifying a species, using ordinary-least-squares regression.

We analysed responses to Q13–Q16 with CLMMs, with pest species, use of citizen science platforms and their interaction as fixed effects, and respondents’ ID as random factor. We used the same modelling approach as described for Q1-Q3.

## Results

### Respondents’ profiles

The survey was opened by 152 persons and completed by 101 respondents. Despite our efforts to balance the number of respondents from the forest and urban sectors, the latter was twice as much more represented **(Table 2)**. Responses originated from 48 administrative regions (so called *départements*, **Figure S1, Supplementary Material**). Slightly more than half of the respondents (57%) had professional activities related to tree, green space or forest management; others declared that their activity involved tree health surveillance. Of those, half were active in the forest sector, and the other half in urban and peri-urban environments **(Figure S2)**.

**Table 2.** Summary of demographic characteristics of respondents.

| **Gender** |  | **Main declared activity** |  |
| --- | --- | --- | --- |
| Male | 62 | Management | 58 |
| Female | 36 | Surveillance | 43 |
| Other | 3 | Sector of activity |  |
| **No years of professional experience with trees** |  |  |  |
| < 5 | 24 | Agriculture | 2 |
|  |  | Urban and peri-urban green spaces | 68 |
| 5-10 | 30 | Forests, urban and peri-urban green species | 11 |
| 10-20 | 25 | Forests | 20 |
|  |  | **Participation in the Surveillance of Regulated and Emerging Organisms (SORE)** |  |
| > 20 | 22 | No | 22 |
| **Highest diploma obtained** |  | Yes | 79 |
| Lower secondary | 4 |  |  |
| Upper secondary (general) | 3 |  |  |
| Upper secondary (vocational) | 8 |  |  |
| Short-cycle tertiary | 25 |  |  |
| Bachelor | 22 |  |  |
| Postgraduate degree («Maîtrise») | 6 |  |  |
| Master | 29 |  |  |
| Doctorate | 1 |  |  |
| Other | 3 |  |  |

Roughly half of respondents (53%) were involved in the official national surveillance scheme for regulated and emerging organisms (i.e., *experts*), to which should be added who declared that their main activity was tree health surveillance, although they were not included in official surveillance schemes at national level (31%). A minority of respondents (22%) declared taking no part in tree health surveillance but only activities related to tree or forest management **(Table 2, Figure S2)**.

### Knowledge of regulated organisms

Differences in the level of knowledge that respondents had of the selected pests and pathogens differed significantly among the species, but this effect was contingent on the respondent status according to their participation in the surveillance of regulated and emerging pests (*i.e.*, significant interaction: LR = 47.08, *P*-value < 0.001). Whereas experts reported similar levels of knowledge across the selected species, those not involved in surveillance reported greater knowledge of the pine and oak processionary moths and of plane canker than of the two *Agrilus* species **(Figure 1)**. Respondents that were not involved in the surveillance had a better knowledge of non-regulated pests in Annex III than of quarantine pests in Annexes II A and II B (Interaction: LR = 37.01, *P*-value < 0.001, **Figure S3**). There was no statistically significant differences in the level of knowledge both groups of respondents had of pests and pathogens (interaction: LR = 0.01, P < 0.001; effect of functional guild: LR = 1.07, P < 0.001).

**Figure 1:**
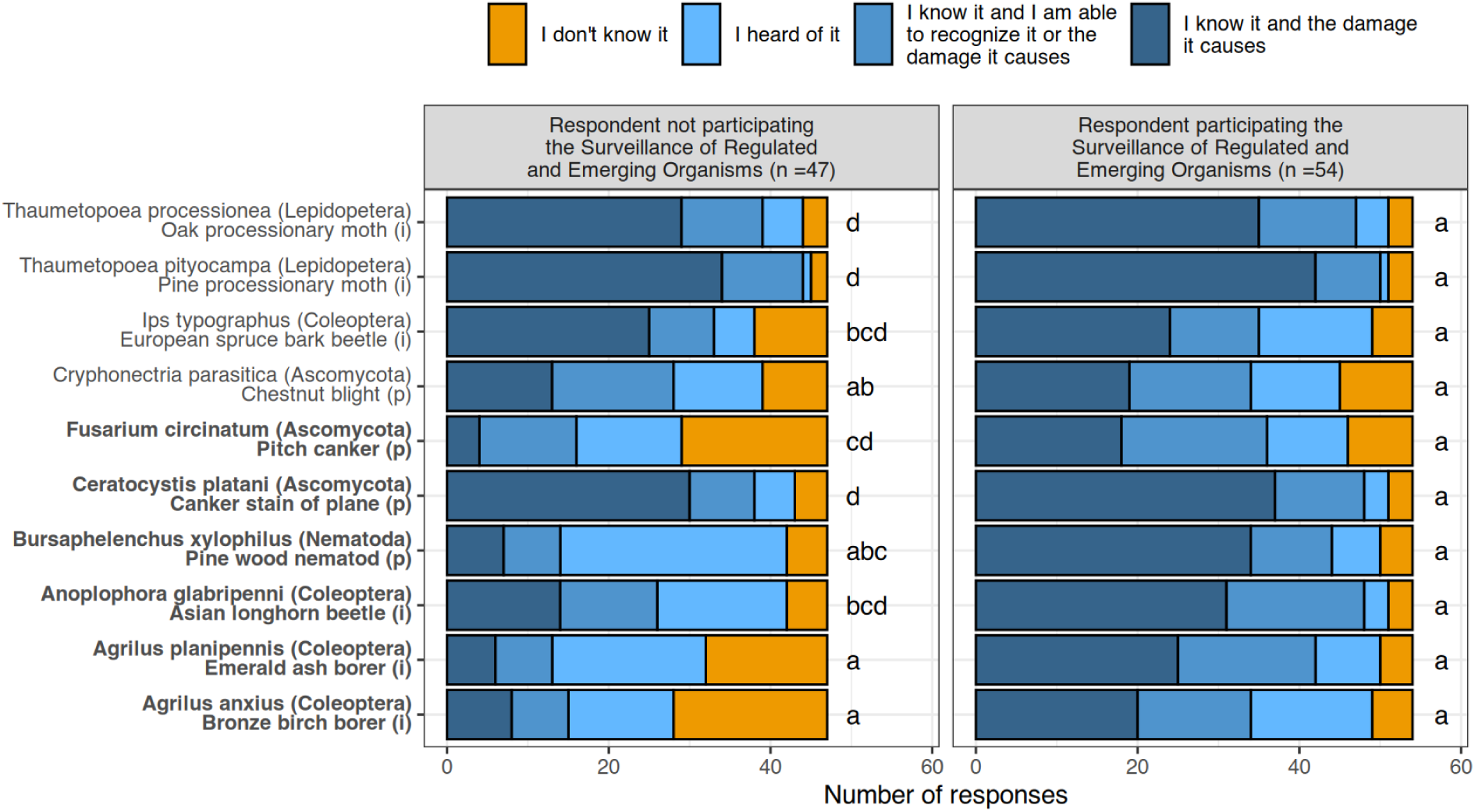
Level of knowledge respondents have of focal pests and pathogens depending on the participation in the surveillance of regulated and emerging pests. Names of pests and pathogens in bold correspond to quarantine species in Annex II A and II B of EU Regulation 2019/2072. (i) and (p) refer to insects and pathogens, respectively.

The number of respondents who correctly assigned quarantine status to regulated organisms and a non-quarantine status to non-regulated organisms differed significantly across species (LR = 188.03, *P*-value = 0.375), and between experts and other respondents (LR = 41.45, *P*-value = 0.375), in a similar way (*i.e.*, no significant interaction: LR = 9.71, *P*-value = 0.375). A greater number of respondents participating in the surveillance assigned a correct status to tested pests and pathogens. Both groups of respondents provided a greater number of correct responses for the plane canker, the pine wood nematode and the Asian longhorn beetle as compared to the two processionary moths, the European spruce bark beetle and the chestnut blight that are all non-regulated organisms **(Figure 2)**. Generally speaking, the number of correct answers was lower for species in Annex III, intermediate for species in Annex II A, and higher for species in Annex II B **(Figure S4)**. The status of insect pests was better known to experts than the status of pathogens, whereas the opposite was true for respondents not involved in surveillance (interaction: LR = 3.37, P = 0.375, **Figure 2**).

**Figure 2:**
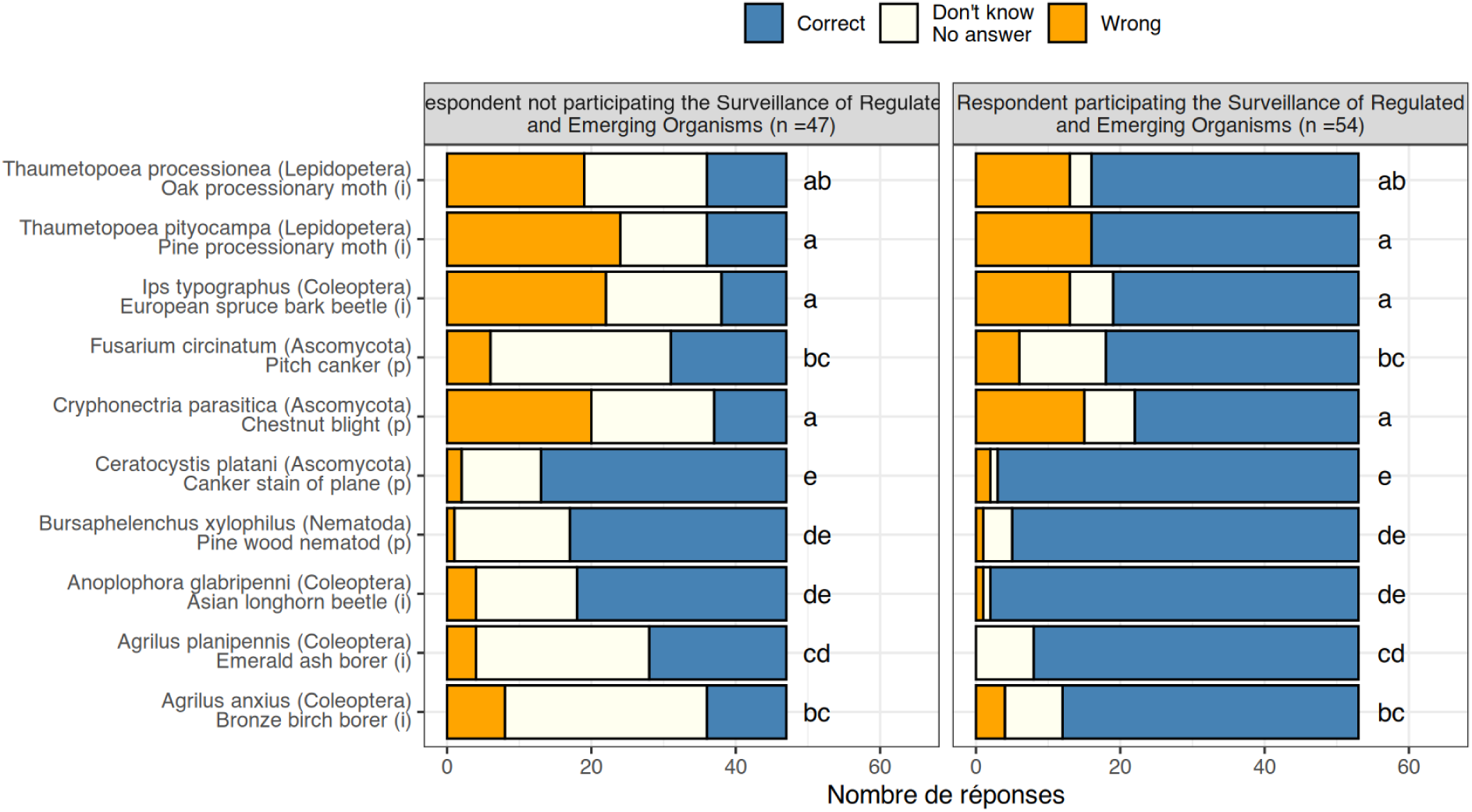
Number of respondents correctly assigning pest to its regulation status (quarantine vs. non-quarantine) depending on their participation in the surveillance of regulated and emerging pests. Names of pests and pathogens in bold correspond to quarantine species. (i) and (p) refer to insects and pathogens, respectively.

### Attitude towards citizen science

Of the 101 respondents, 47 declared having used citizen science platforms (either those we listed or others) and 49 declared using none; 5 did not answer. Among users, 29 were involved in the surveillance of regulated and emerging organisms. Further descriptive analyses focus only on respondents using citizen science platforms. *iNaturalist* was the most frequently used platform, with respondents involved in SORE using it twice as much as other respondents **(Figure S5)**.

Respondents declared using citizen science platforms for both personal and professional use, the latter being more frequent **(Figure 3A)**. When using citizen science platforms in a professional context, 60% of respondents declared doing so on their own initiative **(Figure 3B)**. Only a quarter of respondents declared uploading data to citizen science platforms **(Figure3C)**. Instead, citizen science platforms were used to help with species identification or to visualise the distribution of a particular species **(Figure 3D)**. More than half of the respondents using citizen science platforms in a professional context did so on a monthly basis or more frequently **(Figure 3E)**. We refrained from investigating cross-category interactions because of the overall low number of respondents declaring use of citizen science platforms.

**Figure 3:**
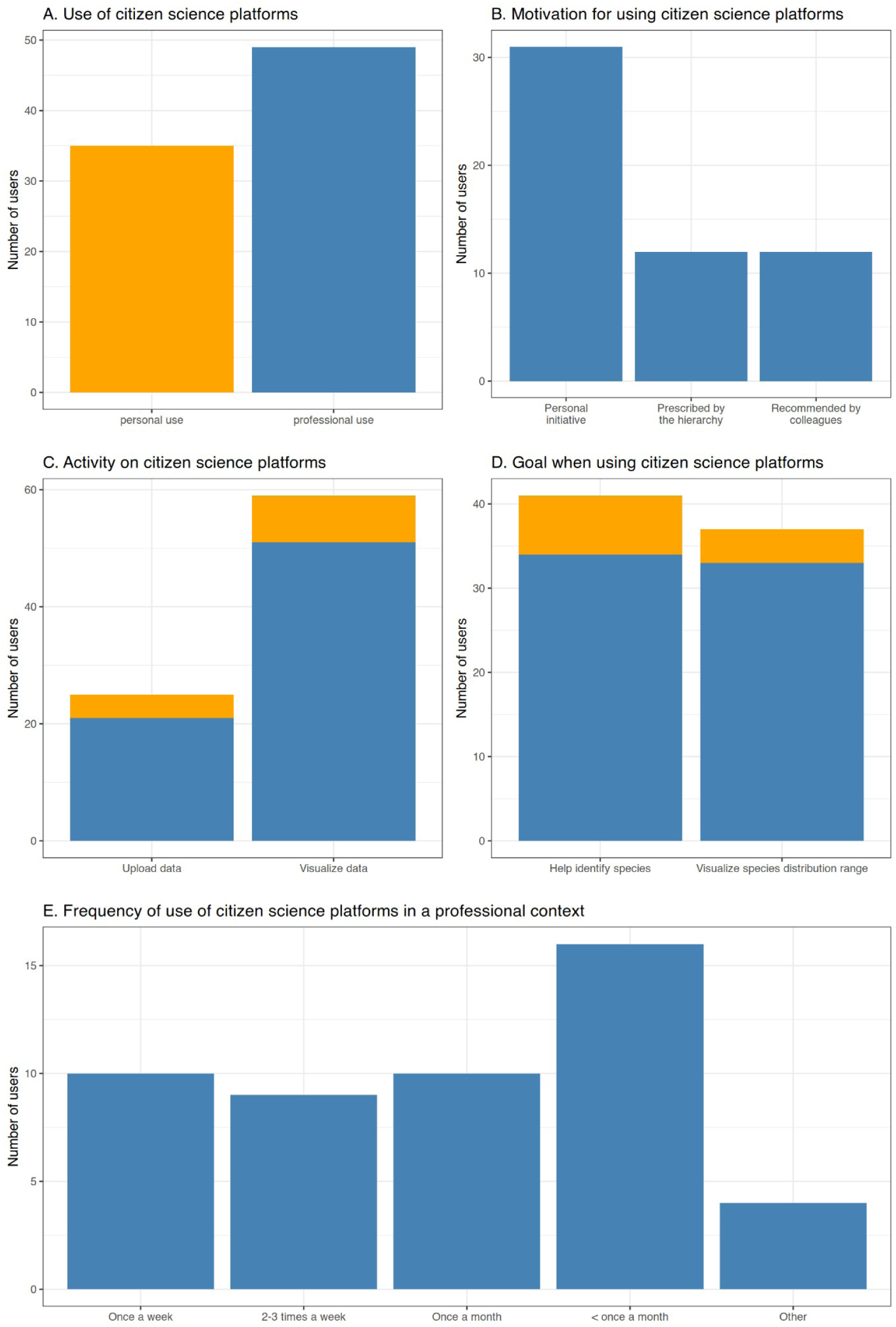
Summary statistics describing in which context (A), why (B), how (C), what for (D) and how often (E) citizen science platforms are used by respondents. Colours in (A) refer to the use of citizen science platforms in a personal (orange) and professional context (blue). The same colours are used in each panel.

### Attitudes towards citizen science platforms as a potential tool for the surveillance of regulated species in France

The two processionary moths and, to a lesser extent, the European spruce bark beetle and the Asian longhorn beetle were considered relatively easy to identify by respondents **(Figure S6)**, and a majority of respondents considered it likely or very likely that they would consult citizen science platforms to obtain information on their presence in a given territory **(Figure S6B)**. However, a substantially smaller proportion of respondents considered that observations of these species would be reported on citizen science platforms **(Figure S6C)**. When considering the tested species as a whole, respondents were more likely to consult citizen-science platforms to obtain information about the presence of a pest or disease when they considered that organism easy to identify (slope ± se: 0.93 ± 0.11, *t*-value = 8.59, *P*-value < 0.001, **Figure 4**). They also regarded occurrences of pests and diseases as being more likely to be reported on citizen-science platforms when the species were easier to identify (0.22 ± 0.05, *t*-value = 4.42, *P*-value = 0.002, but the association between identification and signaling scores was weaker **(Figure 4)**. Respondents with the most experience tended to have a less positive perception of the ease of identifying pests and pathogens and of the likelihood that observations would be reported on citizen science platforms **(Figure S7)**.

**Figure 4:**
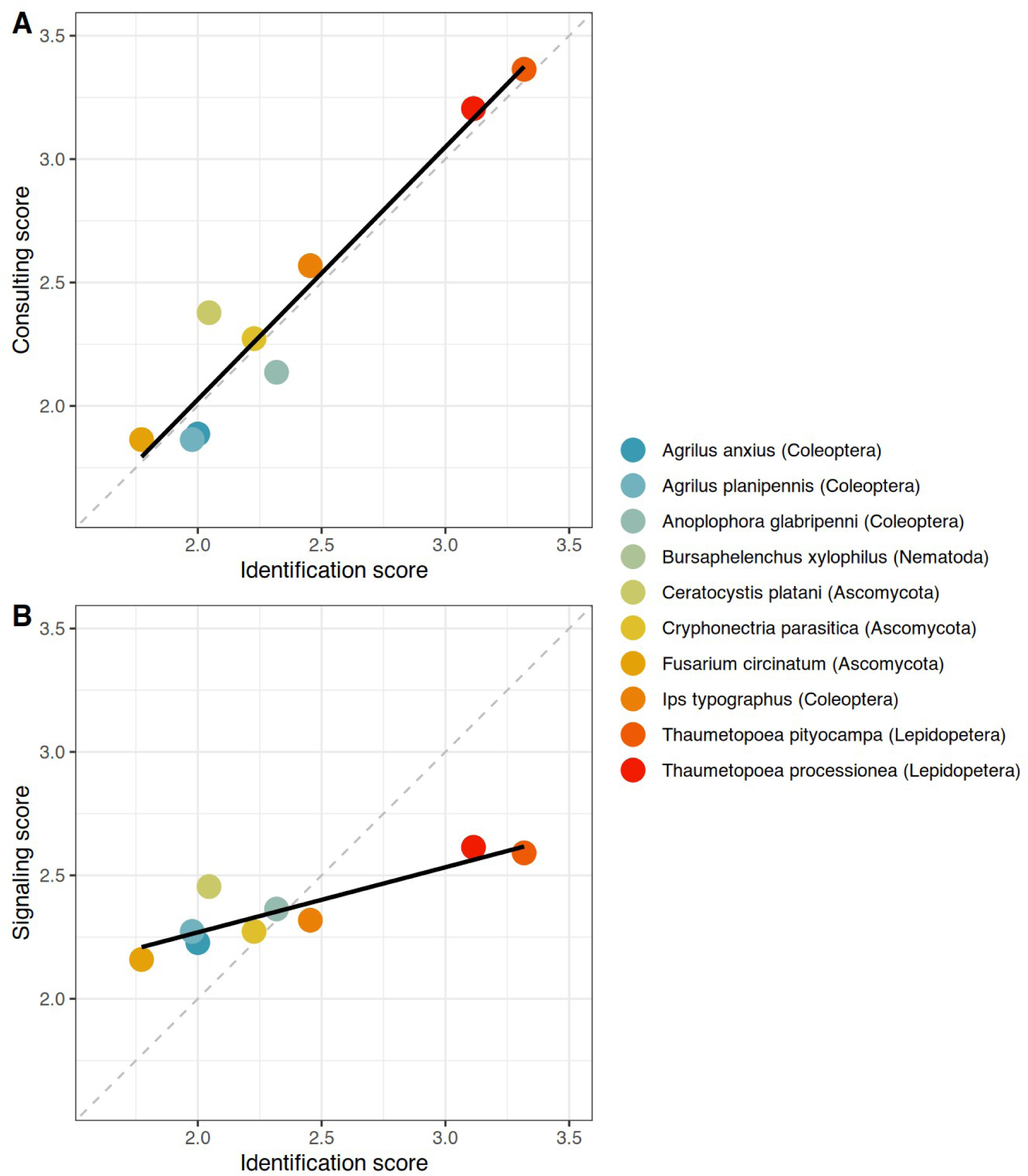
Relationship between respondents’ opinion on the difficulty to identify pests and pathogens and the probability observations are reported on citizen science platforms (signaling) or the probability they would consult these platforms to obtain information regarding the distribution of each pest and pathogen. Dots are averaged scored for each pest or pathogens. The solid line represent the prediction from the ordinary least square regression. The dashed line is the 1:1 relationship.

### Scenarios of use of citizen science platforms

When asked directly, respondents were fairly positive about the potential of using citizen science platforms to aid the early detection of the Asian longhorn beetle *Anoplophora glabripennis* and the oak lace bug *Corythucha arcuata*, with more than three quarters of responses being “agree” or “totally agree”. However, when asked specific questions regarding the information potentially provided by these platforms, their opinions varied among uses (LR = 75.23, *P*-value = 0.57) and between species (LR = 5.93, *P*-value = 0.57), but there was no evidence of significant interaction between both (LR = 2.01, *P*-value = 0.57). Specifically, respondents’ opinions were more positive regarding the potential use of citizen science for the surveillance of *C. arcuata* than that of *A. glabripennis* **(Figure 5)**. They were also more positive about citizen science platforms as a relevant tool to inform on the presence and distribution of these pests, but not to provide reliable information on absence or abundance **(Figure 5)**.

**Figure 5:**
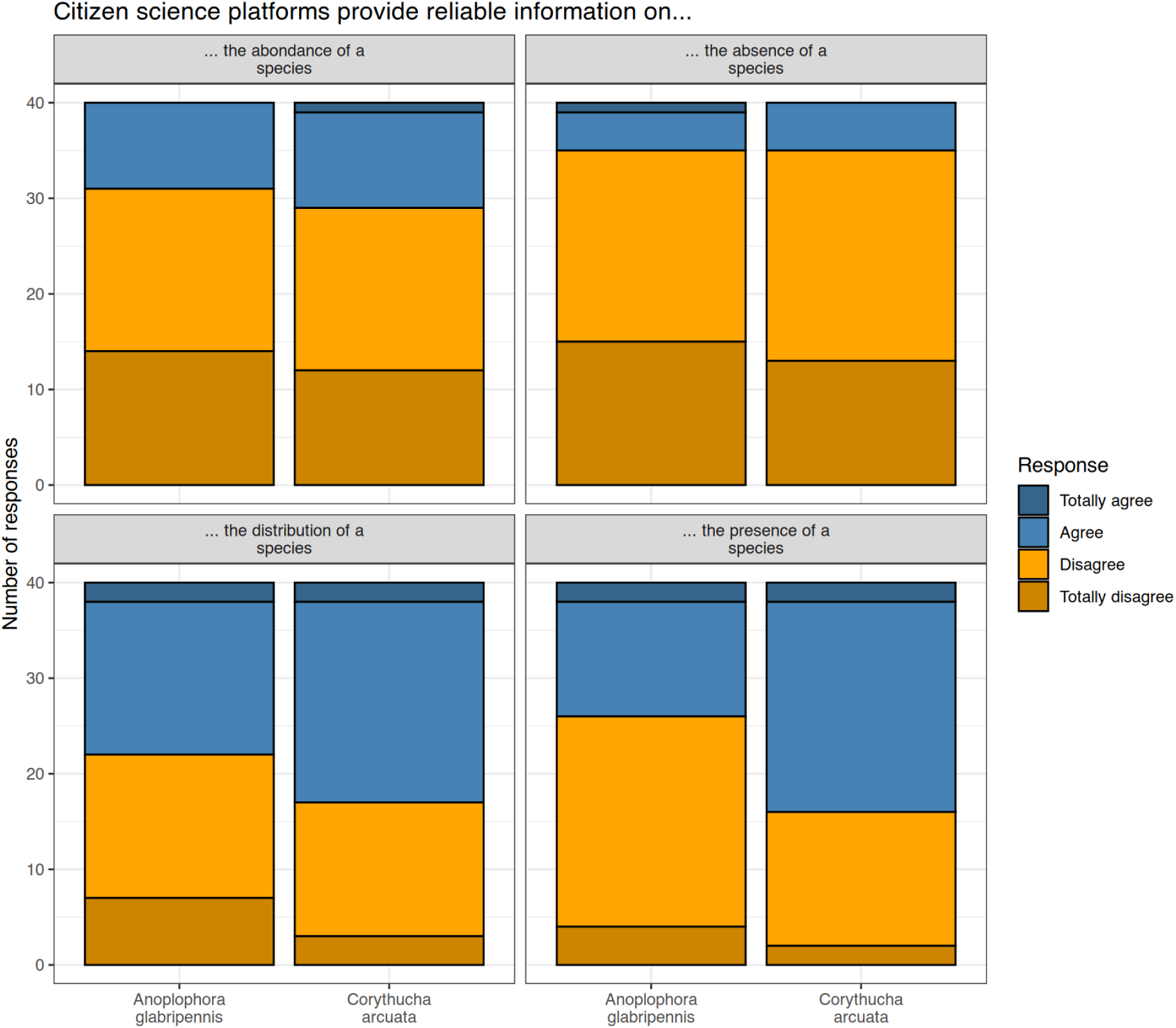
Respondents’ opinion on the potential use of citizen science platforms to obtain reliable information in the presence, absence, distribution and abundance of *Anoplophora glabripennis* and *Corythucha arcuata* in France.

## Discussion

Our survey of stakeholders involved in the management and surveillance of trees and planted, natural and rural forests in France reveals a substantial knowledge of quarantine pests, but mixed attitudes regarding the use and potential of citizen science platforms as relevant sources of information for the early detection and surveillance of regulated tree pests and diseases.

### Tree and forest managers have limited knowledge of regulated pests and pathogens and their EU regulatory status

The survey targeted on the one hand experts appointed to the surveillance of regulated and emerging organisms and on the other hand tree and forest professionals having a range of activities including the management and surveillance of trees, with varying degrees of involvement in tree health surveillance. Experts declared having a good knowledge of both regulated and non-regulated pests, whereas the knowledge of other professionals varied across the tested species.

The surveillance of regulated and emerging pests in France is undertaken by agents employed by different institutions, with sectoral specialisation. For instance, experts in charge of forest health surveillance are not appointed to the surveillance of urban trees or orchards, and *vice versa*. Yet some pests included in our survey are typical forest pests (e.g. pinewood nematode, European spruce bark beetle) whose occurrence in cities is unlikely. Conversely, the canker stain of plane is of greater concern in cities and along roadsides than in forests. Despite this sectoral specialisation, we found no significant difference in the knowledge experts declared having of the different species.

The situation was different for the group of respondents who were not involved in the surveillance of regulated and emerging pests. These respondents declared having a good knowledge of species that widely distributed in France and that are not quarantine species at the EU level (i.e., species listed in Annex III of EU Regulation 2019/2072). However, their knowledge of quarantine pests was lower, particularly for those species that are not known to occur in the European Union (i.e. species listed in Annex IIA). This result is consistent with previous studies revealing that stakeholders’ knowledge of regulated pests is limited, particularly for exotic pests that are not present in the country (Marzano et al. 2016; Raum et al. 2023). Importantly, non-experts had little knowledge of the regulatory status of the set of species we considered in this study; half of them correctly assigned a quarantine status to quarantine species listed in Annex IIB (those known to occur, or to have occurred, in the EU), but had much more limited knowledge of the regulatory status of both non-regulated species (wrongly regarded as regulated) and regulated species that are not present in the EU.

Altogether, these results indicate that current post-border biosecurity procedures could be improved by promoting both knowledge of regulated organisms and awareness of biosecurity schemes (Ram et al. 2016; Warman et al. 2025), at least in France, though questions remain about how to effectively engage stakeholders to promote behavioural change (Hall et al. 2020).

### Citizen science platforms are known, but only used as punctual sources of information

Early detection of regulated pests and pathogens can benefit from citizen scientists reporting observations of plants and animals on online platforms, thus suggesting that citizen science can be a lever for effective biosecurity (Epanchin-Niell et al. 2021; Roe et al. 2024; González-Moreno et al. 2025). However, this would require that, on the one hand, the people most likely to encounter species of concern report their observations and, on the other hand, that these platforms are known to plant protection officers, trusted as relevant sources of information, and ultimately used by them (de Groot et al. 2023). In these respects, we found mixed evidence.

Half of the respondents declared using citizen science platforms, including a majority of experts involved in the surveillance of regulated and emerging pests. Most respondents used citizen science platforms in a professional context, mostly on their own initiative. Interestingly, they declared doing so to help identify species or to track the presence or distribution of a particular species, but none of the respondents using these platforms in a professional context declared sharing their own observations. These results indicate that citizen science, as a means of producing and sharing data, has not yet been integrated into the professional routines of tree and forest professionals, even though half of the respondents reported some interest in data generated through these platforms.

### The perceived relevance of citizen science for biosecurity varies among pests depending on their regulatory status and conspicuousness

Citizen science data are often considered by experts to be unreliable because they suffer from several biases (Caley et al. 2020; de Groot et al. 2022; Pocock et al. 2024), thus limiting their use for post-border biosecurity. Our survey was designed to specifically address the most common beliefs regarding these data, which are frequently regarded as spatially biased (species occurrences indicate where observers go or do not go rather than where species actually occur) (Baker et al. 2019; de Groot et al. 2022) and taxonomically biased, whereby large, conspicuous species are more likely to be reported than small, inconspicuous ones (Caley et al. 2020). Although we found evidence that respondents share these common beliefs, the detail of the responses they provided draws a more nuanced and encouraging picture. Contrary to what would be expected based on their morphology and dangerousness (Caley et al. 2020), respondents were more positive towards the use of citizen science platforms for the detection of the oak lace bug — a grey bug less than 3 mm long — than for the detection of the large, bright Asian longhorn beetle. This difference might be due to the abundance of the former and the conspicuousness of the damage it causes to deciduous oaks. The oak lace bug is now widespread in south-western France, from where it has spread northward and westward over the last few years. It causes clearly visible discolouration in deciduous oaks as early as mid-summer, so that its presence is easily noticed, even by non-experts. By contrast, the Asian longhorn beetle has not established in the country. Although it can drive trees of several species to death, it does not cause clearly distinctive symptoms and does not reach population levels as high as those attained by the oak lace bug, which might explain the difference in stakeholders’ appreciation of the two species. Altogether, these results suggest that stakeholders might consider citizen science platforms more suitable for the monitoring of pests than for the early detection of regulated species (Pocock et al. 2024).

We found that the degree of expertise respondents considered necessary to identify pests and diseases varied among pests, and that it influenced not only their opinion regarding the probability that observations of these pests would be reported on citizen science platforms, but also the probability that they, as stakeholders, would consult these platforms to obtain information on the distribution of pests and diseases. Specifically, we found a strong match between respondents’ opinion on the ease of identifying species and the probability that they would consult citizen science platforms. Interestingly, the species regarded as the easiest to identify are also those best known by respondents. There was also a positive correlation between opinion on the ease of identifying species and the belief regarding the propensity of citizen scientists to report their observations on citizen science platforms.

### Implications for biosecurity

Although we are well aware that these results are based on only a limited number of responses, we see two main implications for tree and forest biosecurity. First, the fact that citizen science platforms are known and used by stakeholders indicates that there is potential for them to be used for the early detection of regulated organisms, as advocated by several authors (Gupta et al. 2022; Hulbert et al. 2023). It also indicates that, despite concerns regarding biases in citizen science data, stakeholders regard these platforms as potentially relevant sources of information. If so, there is a chance that observations of regulated pests shared on citizen science platforms could be noticed by SORE experts and trigger field campaigns aimed at confirming or refuting the observation. By targeting areas worthy of investigation (de Groot et al. 2022), this approach has the potential to accelerate the early detection of newly introduced regulated pests (González-Moreno et al. 2025) and the implementation of appropriate measures. Second, if the former interpretation is valid, encouraging tree and forest professionals to report observations on these platforms, or facilitating the flow of data between citizen science platforms and the databases managed and maintained by plant protection professionals, has the potential to further increase the value of citizen science platforms. We refrain from speculating on the reasons that might explain the observed discrepancy between using citizen science platforms to consult data and using them to contribute data, particularly data derived from the institutional surveillance of regulated pests and pathogens by the competent authorities. We acknowledge this could imply profound changes in agents’ workflows as well as cultural change, which represents a major limiting factor (Magarey et al. 2009; Warman et al. 2025).

### Limitations

Because pests and pathogens can move freely across administrative borders, we intended to survey stakeholders from both the urban and forest sectors, and disseminated the questionnaire to both professional networks. However, after two weeks, we noticed that the vast majority of respondents had professional activities related to trees in urban environments and trees accompanying infrastructure, but far fewer strictly related to forests. We used various approaches to disseminate the questionnaire to stakeholders in the forest sector, including forest managers, advisors, owners, and the network of so-called *Correspondants-Observateurs* who are forest professionals appointed to the surveillance of regulated and emerging pests in France. Despite these efforts, direct contacts, and contacts with their hierarchy, we obtained a limited number of responses from the forest sector. We are aware that this limitation constrains the breadth of our interpretations, but we consider them instructive nonetheless, as they might reveal ambivalent attitudes towards citizen science that would differ between sectors, or a lack of interest in this type of research, which is not directly useful for their purpose.

We cannot rule out that our having initiated the survey only a few months after the pinewood nematode was discovered in south-western France partly explains the lack of responses from the forest sector. Stakeholders involved in the crisis management may indeed have been overwhelmed by these emergencies. However, we note that this would only have concerned experts in the area under scrutiny, and would therefore not explain the lack of responses from stakeholders in other regions. Another possibility is that this crisis has revealed tensions among stakeholders — including authorities, research institutes, forest owners, and forestry companies — that have hampered their willingness to contribute to this research.

## Conclusion

Our survey reveals ambivalent attitudes among tree and forest stakeholders regarding online citizen science platforms and their potential use for the surveillance of tree pests and diseases. First, it highlights that professionals whose activity does not involve the surveillance of regulated and emerging organisms have a limited knowledge of quarantine pests, which could be partly extended to tree biosecurity more broadly. Yet engaging with citizen science has the potential to raise awareness of the risk and its mitigation (**ref?**). Second, citizen science platforms are known to, and used by, half the responding stakeholders, including a majority of experts involved in the surveillance of regulated and emerging pests. However, they are used exclusively as a source of information rather than as a means of sharing information with others. This one-way usage by those most likely to make relevant observations in the field likely slows the diffusion of knowledge and awareness of the risk within the wider community. Finally, our results allow us to infer that professionals are note reluctant to consider citizen science platforms as a companion tool for tree biosecurity experts, helping to survey the spread of easily identifiable pests. However, we found no indication that citizen science can readily be integrated as a routine tool for the early detection of quarantine pests in France without further methodological and operational developments, the feasibility of which warrants further investigation.

## Acknowledgements

This research has received funding from the European Union’s Horizon Europe Research and Innovation programme under grant agreement No. 101134200 “FORSAID: Forest surveillance with artificial intelligence and digital technologies”. A CC-BY public copyright license has been applied by the authors to the present document and will be applied to all subsequent versions up to the Author Accepted Manuscript arising from this submission, in accordance with the grant’s open access conditions. The authors are grateful to Maxime Guérin from the Plante & Cité association and the members of the BIODIV team at the BIOGECO laboratory (INRAE, University of Bordeaux) for testing the preliminary versions of the questionnaire and for their insightful comments on the wording of the questions and response categories. The authors declared having used AI assisted copy editing to improve fully human-generated texts to ensure that the texts are free of errors in grammar and spelling.

## Statements & Declarations

## Authors contributions

BC and MdG initiated the project. All authors developed the concept of the study. BC, SB, AM and BB designed the questionnaire. SB prepared and administrated the questionnaire. BC analysed the data with conceptual inputs from all authors. BC drafted the first version of manuscript. All authors contributed critical comments and edits.

## Data availability

The dataset generated and analysed during the current study is available in the https://recherche.data.gouv.fr/fr; [**ADD-LINK**]

## Consent to participate

Informed consent was obtained from all individual participants included in the study as part of the questionnaire.

