## Supplementary Material for "Knowledge gap and conditional receptivity: why tree and forest professionals underutilize citizen science for quarantine pest detection in France"

### A survey of professional knowledge and attitudes towards citizen science for tree pest surveillance in France

—

### Supplementary tables and figures

Table S1. List of agency to which the survey was distribute at the initial step of the snowball approach.

| Sector | Agency | Type of agency |
|---|---|---|
| Forest | DSF | Public |
| | ONF | Public |
| | CNPF, CRPF | Public |
| | IGN | Public |
| | Alliance Forêt Bois | Private |
| | Association des entrepreneurs des travaux forestiers | Private |
| | Les experts forestiers | Private |
| | Canopée | Private |
| Urban gardens, green spaces and green infrastructures (JEVI) | Référent JEVI | Public, Private |
| | Bordeaux métropole | Public |
| | Association française d’arboriculture | Private |
| | Ecole nationale du paysage | Private |
| | Plante et cité | Private |
| | Fédération française du paysage | Private |
| | SEQUOIA | Private |
| | Groupement des experts conseils en arboriculture ornementale | Private |
| | Union nationale des entreprises du paysage (UNEP) | Private |
| | VALHOR | Private |
| | POLLENIZ | Private |

| | | |
|---|---|---|
| | CDHR | Private |
| Forest & JEVI | Fredon France | Public, Private |
| | Fredon Nouvelle Aquitaine | Public, Private |
| | Fredon Nantes | Public, Private |

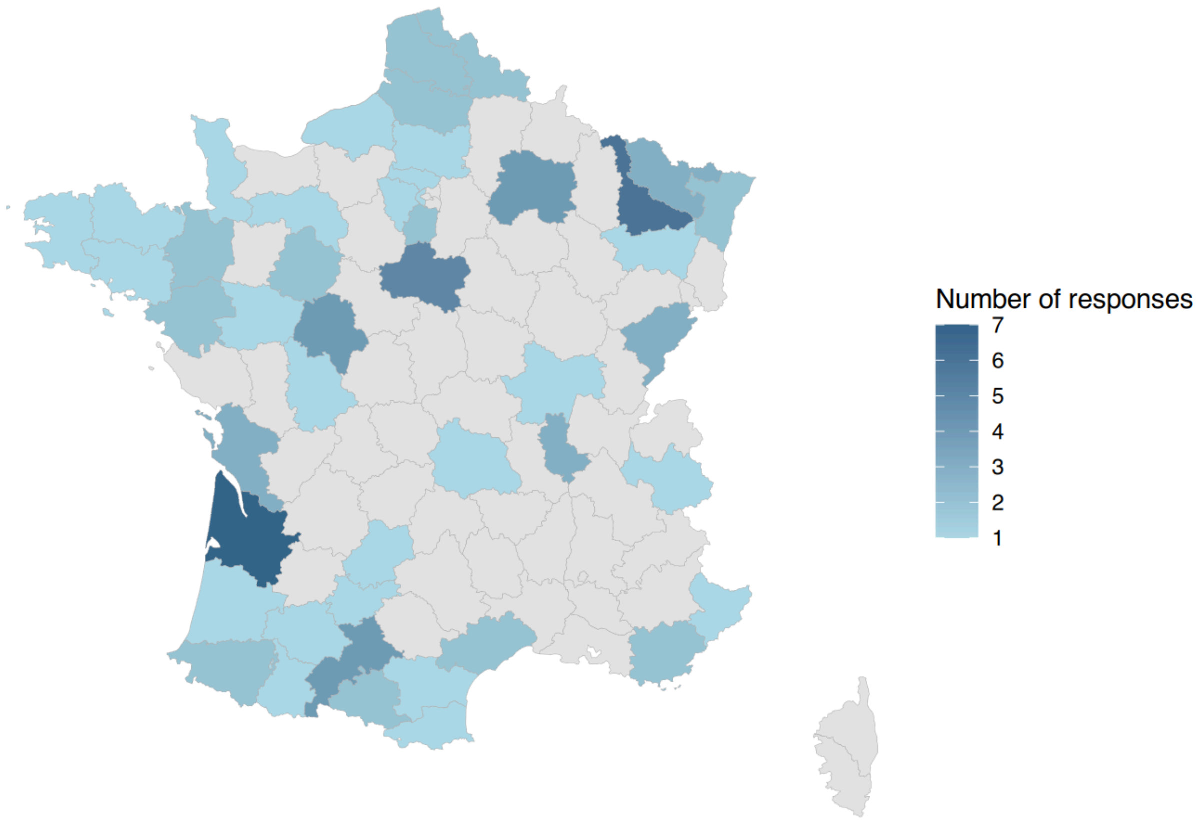


**Figure S1:** Origin of responses spilt according to French administrative regions.

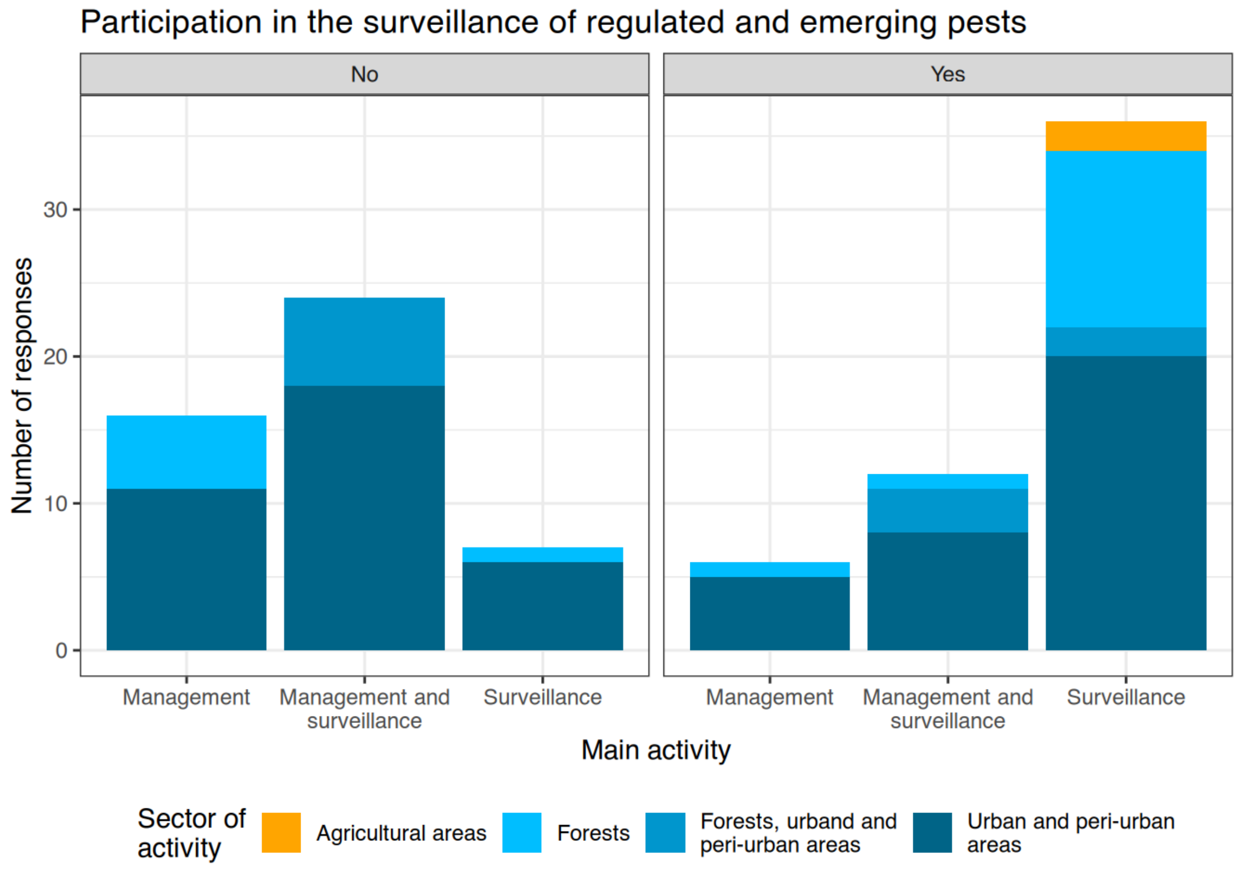


**Figure S2:** Number of respondents whose main activity is related to tree surveillance, management or both, depending on their sector of activity and their participation to the surveillance of regulated and emerging organisms.

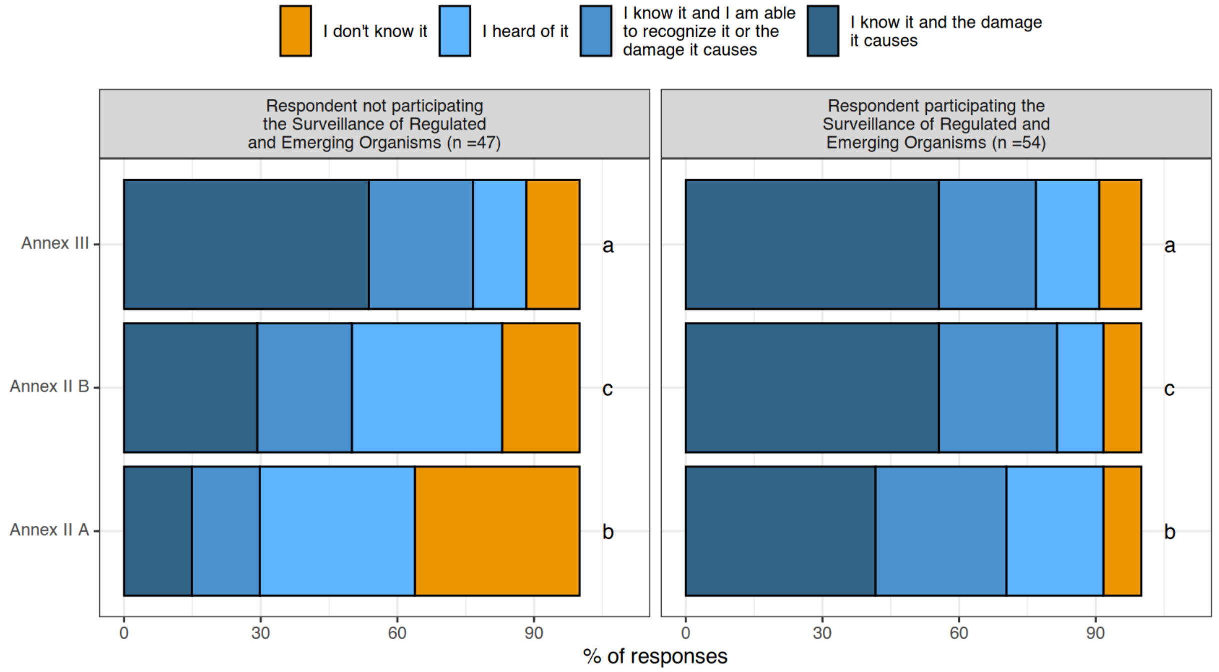


Figure S3: Level of knowledge respondents have of focal pests and pathogens depending on the participation in the surveillance of regulated and emerging pests. (i) and (p) refer to insects and pathogens, respectively. Species listed in Annexes IIA and IIB are EU quarantine species, currently absent from the EU territory (IIA), or known to occur within the EU (IIB). Species listed in Annex III are regulated organisms that are not considered EU quarantine species.

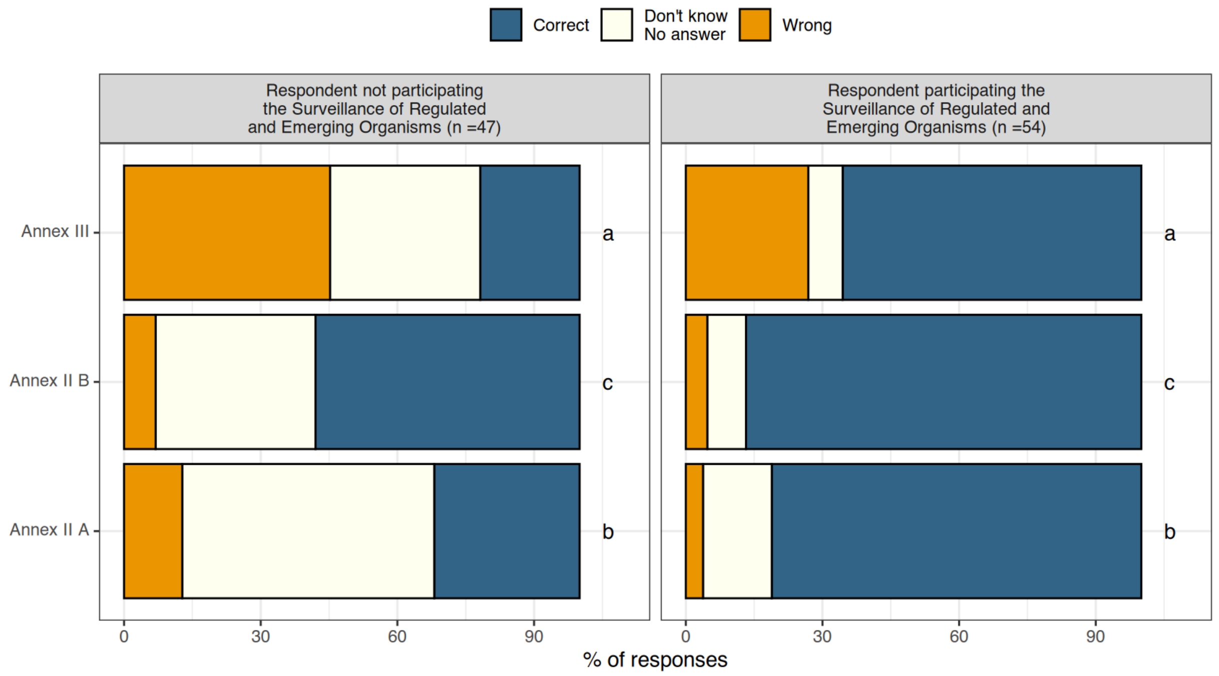


Figure S4: Level of knowledge respondents have of focal pests and pathogens depending on the participation in the surveillance of regulated and emerging pests. (i) and (p) refer to insects and pathogens, respectively. Species listed in Annexes IIA and IIB are EU quarantine species, currently absent from the EU territory (IIA), or known to occur within the EU (IIB). Species listed in Annex III are regulated organisms that are not considered EU quarantine species.

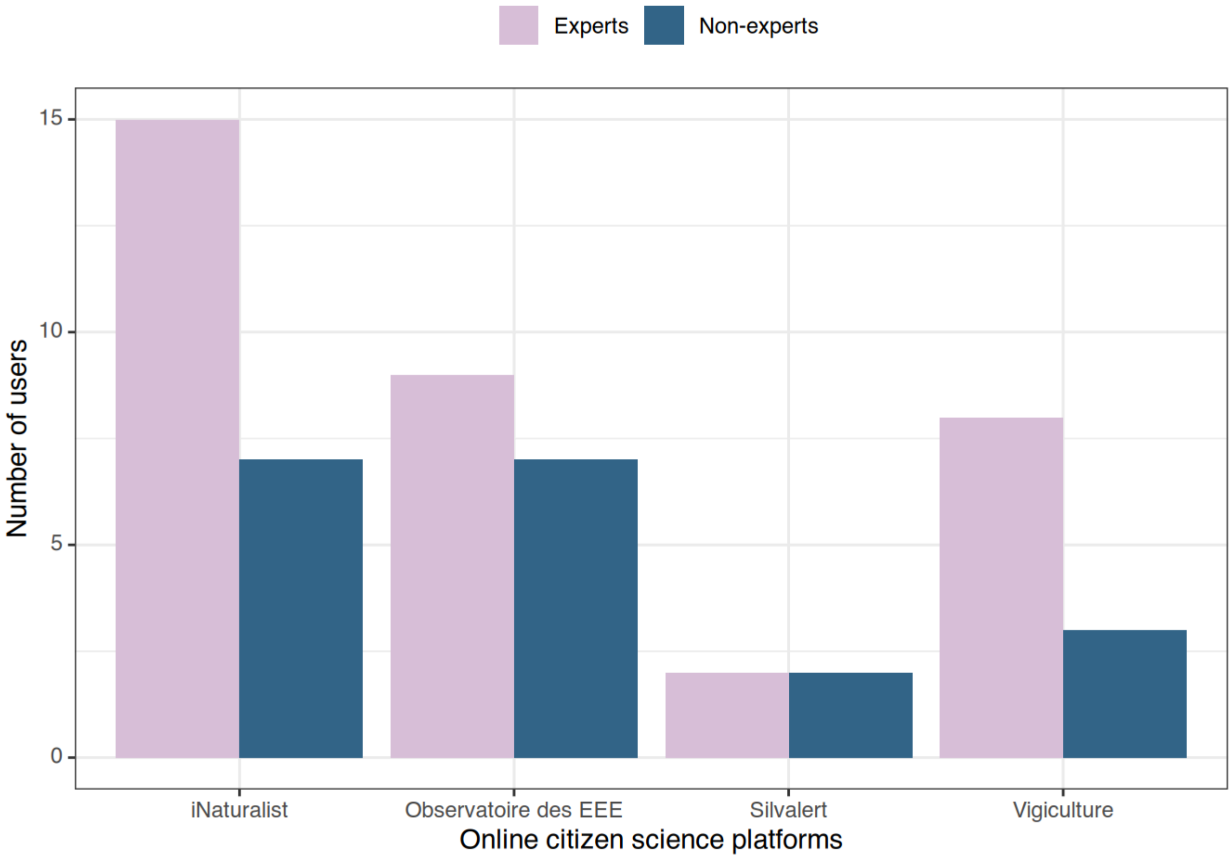


Figure S5: Number of respondents using different citizen science platforms according to their participation in the surveillance of regulated and emerging organisms (SORE).

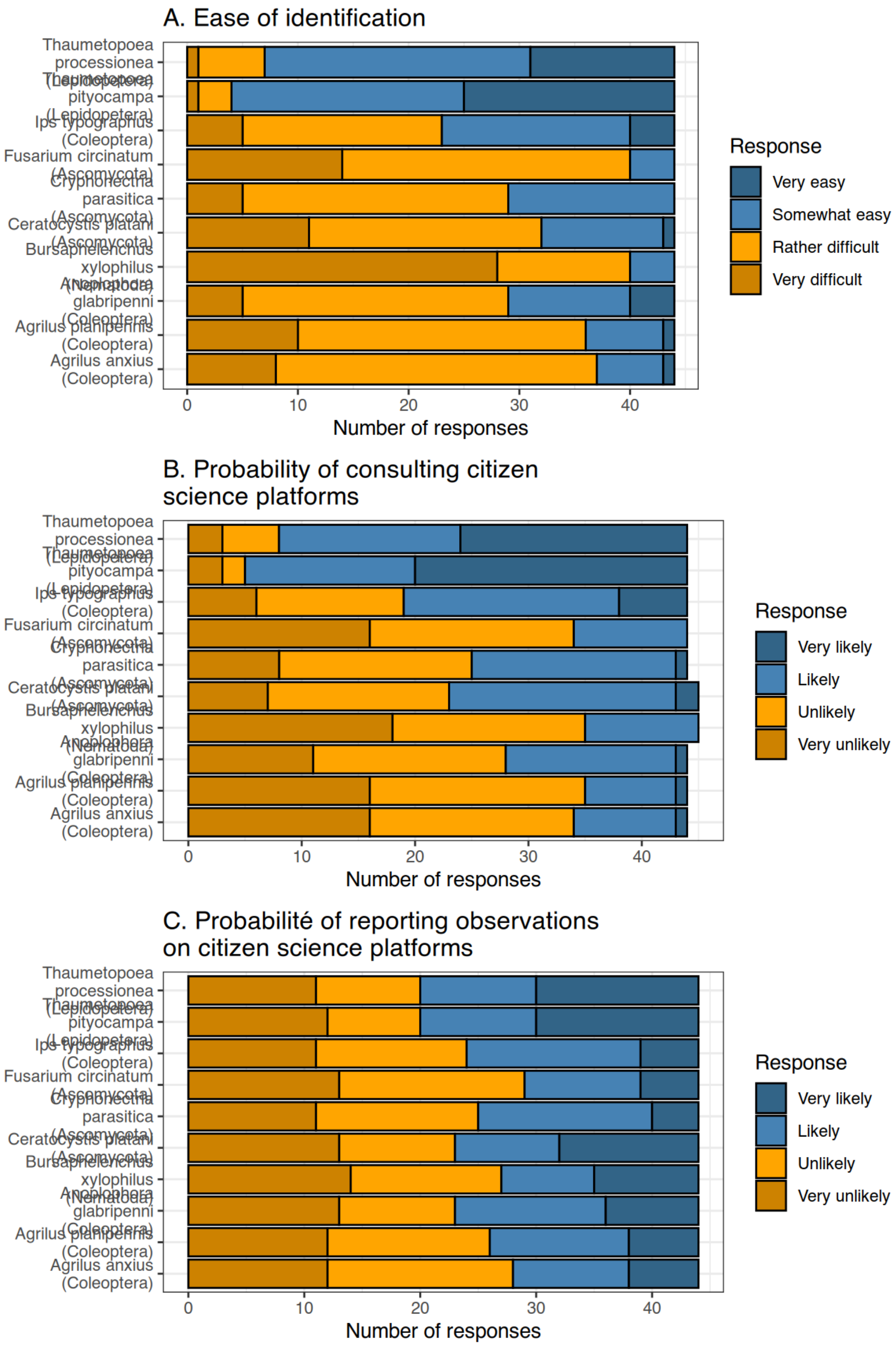


Figure S6: Respondents' opinion on (A) the difficulty of identifying the focal tree pests and diseases, on (B) the probability that users of citizen science platforms would share observations of these species and (C) on the probability that they would consult citizen science platforms to obtain information regarding the distribution of each pest and pathogen.

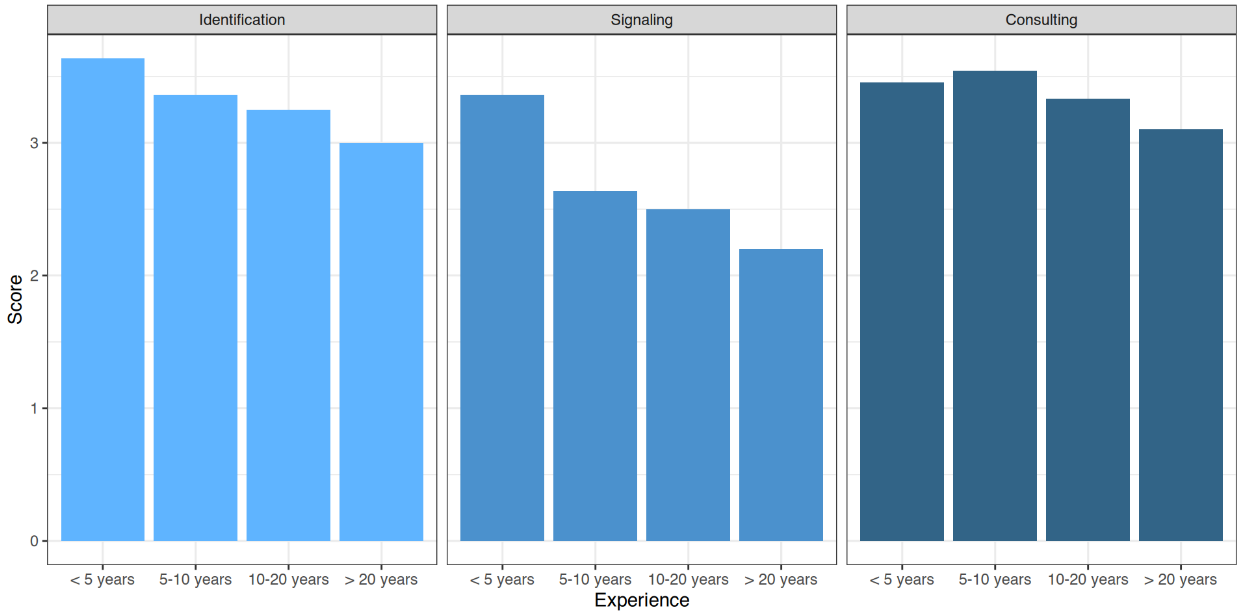

Figure S7: Relationship between the number of years of experience and respondents' opinions regarding the difficulty of identifying pests and pathogens, the likelihood that observations are reported on citizen science platforms (signaling), and the likelihood that they would consult these platforms to obtain information on the distribution of each pest and pathogen.
