## Appendix A for "Knowledge gap and conditional receptivity: why tree and forest professionals underutilize citizen science for quarantine pest detection in France"

Appendix 1 - Survey on the use of citizen science in tree and forest health surveillance

### Survey on the use of citizen science in tree and forest health surveillance

You are invited to take part in a survey aimed at understanding the potential role of citizen science in tree and forest health surveillance. We are particularly interested in your use of digital biodiversity data-sharing platforms in the course of your professional activity.

 Your participation in this questionnaire will help to improve knowledge of the potential role of citizen science in the early detection of quarantine and regulated pests.

#### PROFESSIONAL ACTIVITY

##### Have you worked, or do you currently work, in a professional activity related to trees (in towns, green spaces, forests, etc.)?  *

***Please select only one of the following options:***

- Yes
- No

#### RESPONDENT PROFILE

##### What is your gender?

Answer this question only if the following conditions are met:

The answer was 'Yes' to question '1 [Q1]' (Have you worked, or do you currently work, in a professional activity related to trees (in towns, green spaces, forests, etc.)?)

***Please select one answer below*** :

- Man
- Woman
- Other

##### How old are you?

Answer this question only if the following conditions are met:

The answer was 'Yes' to question '1 [Q1]' (Have you worked, or do you currently work, in a professional activity related to trees (in towns, green spaces, forests, etc.)?)

***Please select one answer below :***

- Under 30
- 30-39
- 40-49
- 50-59
- 60 and over

**What is the highest qualification you have obtained ?**

Answer this question only if the following conditions are met:

The answer was 'Yes' to question '1 [Q1]' (Have you worked, or do you currently work, in a professional activity related to trees (in towns, green spaces, forests, etc.)?)

***Please select one answer below :***

- BEPC (lower secondary certificate)
- Vocational baccalaureate
- General baccalaureate
- BTS (two-year higher technical diploma)
- Bachelor's degree or equivalent (3 years post-baccalaureate)
- Maîtrise or equivalent (4 years post-baccalaureate)
- Master's degree or equivalent (5 years post-baccalaureate)
- Doctorate
- Other

##### What is your job title?

Answer this question only if the following conditions are met:

The answer was 'Yes' to question '1 [Q1]' (Have you worked, or do you currently work, in a professional activity related to trees (in towns, green spaces, forests, etc.)?)

***Tick the answer(s) that apply :***

- Observer Correspondent (DSF, Forest Health Department)
- Officer of the State decentralised services (DRAAF/SRAL)
- Plant health inspector or project officer (DRAAF/SRAL)
- Inspector in the FREDON network
- Plant health adviser
- Phytosanitary inspector
- Forest manager, ONF (National Forests Office)
- Private-sector forest manager
- Forest worker
- Forestry expert
- Forestry adviser (CNPF, CRPF)
- Head of green space services (local authority or private estate)
- Climbing arborist (tree pruner)
- Urban tree heritage manager
- Gardener, green space officer or technician
- Head of green space departments
- Landscape designer
- Forest owner
- FREDON adviser
- Expert and/or adviser in ornamental arboriculture
- Other:

##### Does your professional activity involve:

Answer this question only if the following conditions are met:

The answer was 'Yes' to question '1 [Q1]' (Have you worked, or do you currently work, in a professional activity related to trees (in towns, green spaces, forests, etc.)?)

***Tick the answer(s) that apply :***

- Tree health surveillance (in towns, green spaces, forests, etc.)
- Tree management (in towns, green spaces, forests, etc.)

**Your main area of work concerns:**

Answer this question only if the following conditions are met:

The answer was 'Yes' to question '1 [Q1]' (Have you worked, or do you currently work, in a professional activity related to trees (in towns, green spaces, forests, etc.)? )

***Tick the answer(s) that apply :***

- Forests (planted, natural or semi-natural)
- Urban and peri-urban green spaces (municipal, private)
- Other:

**Number of years of experience in this activity:**

Answer this question only if the following conditions are met:

The answer was 'Yes' to question '1 [Q1]' (Have you worked, or do you currently work, in a professional activity related to trees (in towns, green spaces, forests, etc.)? )

***Please select one answer below :***

- Less than 5 years
- 5 to 10 years
- 10 to 20 years
- More than 20 years

##### Please give the code of the French département in which you mainly carry out your professional activity:  *

Answer this question only if the following conditions are met:

The answer was 'Yes' to question '1 [Q1]' (Have you worked, or do you currently work, in a professional activity related to trees (in towns, green spaces, forests, etc.)? )

***Please write your answer here:………………………………………………………….***

##### Do you take part in the Surveillance of Regulated and Emerging Pests (SORE)?

Answer this question only if the following conditions are met:

The answer was 'Yes' to question '1 [Q1]' (Have you worked, or do you currently work, in a professional activity related to trees (in towns, green spaces, forests, etc.)? )

***Please select only one of the following options:***

- Yes
- No

#### KNOWLEDGE OF REGULATED AND EMERGING PESTS

**Below is a list of tree pests and diseases. For each of them, please tick in this table the statement that best matches your current level of knowledge:**

Answer this question only if the following conditions are met:

The answer was 'Yes' to question '1 [Q1]' (Have you worked, or do you currently work, in a professional activity related to trees (in towns, green spaces, forests, etc.)? )

|  | I am not familiar with this pest/disease | I have already heard of it | I would be able to recognise it and its symptoms | I know it and the damage it causes |
| --- | --- | --- | --- | --- |
| Pine wood nematode *(Bursaphelenchus xylophilus)* |  |  |  |  |
| Emerald ash borer *(Agrilus planipennis)* |  |  |  |  |
| Pitch canker *(Fusarium circinatum)* |  |  |  |  |
| Chestnut blight *(Cryphonectria parasitica)* |  |  |  |  |
| Oak processionary moth *(Thaumetopoea processionea)* |  |  |  |  |
| European spruce bark beetle *(Ips typographus)* |  |  |  |  |
| Bronze birch borer *(Agrilus anxius)* |  |  |  |  |
| Pine processionary moth *(Thaumetopoea pityocampa)* |  |  |  |  |
| Canker stain of plane *(Ceratocystis platani)* |  |  |  |  |
| Asian longhorned beetle *(Anoplophora glabripennis)* |  |  |  |  |

##### Do you think these tree pests or diseases are subject to the Surveillance of Regulated and Emerging Pests (SORE) ? *

Answer this question only if the following conditions are met:

The answer was 'Yes' to question '1 [Q1]' (Have you worked, or do you currently work, in a professional activity related to trees (in towns, green spaces, forests, etc.)?

|  | Yes | No | Don't know |
| --- | --- | --- | --- |
| Pine wood nematode *(Bursaphelenchus xylophilus)* |  |  |  |
| Emerald ash borer *(Agrilus planipennis)* |  |  |  |
| Pitch canker *(Fusarium circinatum)* |  |  |  |
| Chestnut blight *(Cryphonectria parasitica)* |  |  |  |
| Oak processionary moth *(Thaumetopoea processionea)* |  |  |  |
| European spruce bark beetle *(Ips typographus)* |  |  |  |
| Bronze birch borer *(Agrilus anxius)* |  |  |  |
| Pine processionary moth *(Thaumetopoea pityocampa)* |  |  |  |
| Canker stain of plane *(Ceratocystis platani)* |  |  |  |
| Asian longhorned beetle *(Anoplophora glabripennis)* |  |  |  |

##### In your view, the potential presence of these tree pests or diseases in France is:

Answer this question only if the following conditions are met:

The answer was 'Yes' to question '1 [Q1]' (Have you worked, or do you currently work, in a professional activity related to trees (in towns, green spaces, forests, etc.)? )

|  | Not at all worrying | Slightly worrying | Worrying | Very worrying | Don't know |
| --- | --- | --- | --- | --- | --- |
| Pine wood nematode *(Bursaphelenchus xylophilus)* |  |  |  |  |  |
| Emerald ash borer *(Agrilus planipennis)* |  |  |  |  |  |
| Pitch canker *(Fusarium circinatum)* |  |  |  |  |  |
| Chestnut blight *(Cryphonectria parasitica)* |  |  |  |  |  |
| Oak processionary moth *(Thaumetopoea processionea)* |  |  |  |  |  |
| European spruce bark beetle *(Ips typographus)* |  |  |  |  |  |
| Bronze birch borer *(Agrilus anxius)* |  |  |  |  |  |
| Pine processionary moth *(Thaumetopoea pityocampa)* |  |  |  |  |  |
| Canker stain of plane *(Ceratocystis platani)* |  |  |  |  |  |
| Asian longhorned beetle *(Anoplophora glabripennis)* |  |  |  |  |  |

#### USE OF DIGITAL BIODIVERSITY DATA-SHARING PLATFORMS

##### Which of the following digital biodiversity platforms do you use?

Answer this question only if the following conditions are met:

The answer was 'Yes' to question '1 [Q1]' (Have you worked, or do you currently work, in a professional activity related to trees (in towns, green spaces, forests, etc.)? )

***Tick the answer(s) that apply :***

- I do not use any digital biodiversity platform
- iNaturalist
- Observatoire des espèces exotiques envahissantes
- Silvalert
- Vigil’encre
- Vigicultures
- I use other platform(s) not mentioned in this list

##### In which context(s) do you use Vigil’encre?

Answer this question only if the following conditions are met:

The answer was 'Yes' to question '1 [Q1]' (Have you worked, or do you currently work, in a professional activity related to trees (in towns, green spaces, forests, etc.)? ) *and* The answer was 'Vigil’encre' to question '14 [Q14]' (Which of the following digital biodiversity platforms do you use?   )

***Tick the answer(s) that apply :***

- I use it in a personal capacity (walks in nature, taking part in a community project, taking part in a family or friends project, etc.)
- I use it in a professional context (before going into the field, in the field during inspections, during data collection and processing, etc.)

##### Is your use of Vigil’encre in a professional context :

Answer this question only if the following conditions are met:

The answer was 'Yes' to question '1 [Q1]' (Have you worked, or do you currently work, in a professional activity related to trees (in towns, green spaces, forests, etc.)? ) *and* The answer was 'Vigil’encre' to question '14 [Q14]' (Which of the following digital biodiversity platforms do you use?   ) *and* The answer was 'I use it in a professional context (before going into the field, in the field during inspections, during data collection and processing, etc.) ' to question '15 [Q15]' (In which context(s) do you use Vigil’encre? )

***Tick the answer(s) that apply :***

- Required by your management
- Recommended by colleagues
- Recommended by friends/family members
- On your own initiative
- Other:

##### On average, how often do you use Vigil’encre  in a professional context ?

Answer this question only if the following conditions are met:

The answer was 'Yes' to question '1 [Q1]' (Have you worked, or do you currently work, in a professional activity related to trees (in towns, green spaces, forests, etc.)? ) *and* The answer was 'Vigil’encre' to question '14 [Q14]' (Which of the following digital biodiversity platforms do you use?   ) *and* The answer was 'I use it in a professional context (before going into the field, in the field during inspections, during data collection and processing, etc.) ' to question '15 [Q15]' (In which context(s) do you use Vigil’encre? )

***Please select one answer below :***

- Every day
- 2 or 3 times a week
- Once a week
- Once a month
- Less than once a month
- Other

##### What type of use do you make of the Vigil’encre platform?

Answer this question only if the following conditions are met:

The answer was 'Yes' to question '1 [Q1]' (Have you worked, or do you currently work, in a professional activity related to trees (in towns, green spaces, forests, etc.)? ) *and* The answer was 'Vigil’encre' to question '14 [Q14]' (Which of the following digital biodiversity platforms do you use?   )

***Tick the answer(s) that apply :***

- I contribute to the platform with my own observations (sharing data on the platform)
- I consult the data (following observations or reports, collecting data)

##### For what purpose do you use Vigil’encre?

Answer this question only if the following conditions are met:

The answer was 'Yes' to question '1 [Q1]' (Have you worked, or do you currently work, in a professional activity related to trees (in towns, green spaces, forests, etc.)? ) *and* The answer was 'Vigil’encre' to question '14 [Q14]' (Which of the following digital biodiversity platforms do you use?   )

***Tick the answer(s) that apply :***

- Identifying species
- Reporting and sharing observations with other users
- Keeping a record of my observations
- Monitoring the presence of pests in a given area
- Other:

##### In which context(s) do you use Silvalert ?

Answer this question only if the following conditions are met:

The answer was 'Yes' to question '1 [Q1]' (Have you worked, or do you currently work, in a professional activity related to trees (in towns, green spaces, forests, etc.)? ) *and* The answer was 'Silvalert' to question '14 [Q14]' (Which of the following digital biodiversity platforms do you use?   )

***Tick the answer(s) that apply :***

- I use it in a personal capacity (walks in nature, taking part in a community project, taking part in a family or friends project, etc.)
- I use it in a professional context (before going into the field, in the field during inspections, during data collection and processing, etc.)

##### Is your use of Silvalert in a professional context :

Answer this question only if the following conditions are met:

The answer was 'Yes' to question '1 [Q1]' (Have you worked, or do you currently work, in a professional activity related to trees (in towns, green spaces, forests, etc.)? ) *and* The answer was 'Silvalert' to question '14 [Q14]' (Which of the following digital biodiversity platforms do you use?   ) *and* The answer was 'I use it in a professional context (before going into the field, in the field during inspections, during data collection and processing, etc.) ' to question '20 [Q16]' (In which context(s) do you use Silvalert ? )

***Tick the answer(s) that apply :***

- Required by your management
- Recommended by colleagues
- Recommended by friends/family members
- On your own initiative
- Other:

##### On average, how often do you use Silvalert  in a professional context ?

Answer this question only if the following conditions are met:

The answer was 'Yes' to question '1 [Q1]' (Have you worked, or do you currently work, in a professional activity related to trees (in towns, green spaces, forests, etc.)? ) *and* The answer was 'Silvalert' to question '14 [Q14]' (Which of the following digital biodiversity platforms do you use?   ) *and* The answer was 'I use it in a professional context (before going into the field, in the field during inspections, during data collection and processing, etc.) ' to question '20 [Q16]' (In which context(s) do you use Silvalert ? )

***Please select one answer below :***

- Every day
- 2 or 3 times a week
- Once a week
- Once a month
- Less than once a month
- Other

##### What type of use do you make of the Silvalert platform?

Answer this question only if the following conditions are met:

The answer was 'Yes' to question '1 [Q1]' (Have you worked, or do you currently work, in a professional activity related to trees (in towns, green spaces, forests, etc.)? ) *and* The answer was 'Silvalert' to question '14 [Q14]' (Which of the following digital biodiversity platforms do you use?   )

***Tick the answer(s) that apply :***

- I contribute to the platform with my own observations (sharing data on the platform)
- I consult the data (following observations or reports, collecting data)

##### For what purpose do you use Silvalert?

Answer this question only if the following conditions are met:

The answer was 'Yes' to question '1 [Q1]' (Have you worked, or do you currently work, in a professional activity related to trees (in towns, green spaces, forests, etc.)? ) *and* The answer was 'Silvalert' to question '14 [Q14]' (Which of the following digital biodiversity platforms do you use?   )

***Tick the answer(s) that apply :***

- Identifying species
- Reporting and sharing observations with other users
- Keeping a record of my observations
- Monitoring the presence of pests in a given area
- Other:

##### In which context(s) do you use the Observatoire des espèces exotiques envahissantes ?

Answer this question only if the following conditions are met:

The answer was 'Yes' to question '1 [Q1]' (Have you worked, or do you currently work, in a professional activity related to trees (in towns, green spaces, forests, etc.)? ) *and* The answer was 'Observatoire des espèces exotiques envahissantes' to question '14 [Q14]' (Which of the following digital biodiversity platforms do you use?   )

Tick the answer(s) that apply :

- I use it in a personal capacity (walks in nature, taking part in a community project, taking part in a family or friends project, etc.)
- I use it in a professional context (before going into the field, in the field during inspections, during data collection and processing, etc.)

##### Is your use of the Observatoire des espèces exotiques envahissantes in a professional context :

Answer this question only if the following conditions are met:

The answer was 'Yes' to question '1 [Q1]' (Have you worked, or do you currently work, in a professional activity related to trees (in towns, green spaces, forests, etc.)? ) *and* The answer was 'Observatoire des espèces exotiques envahissantes' to question '14 [Q14]' (Which of the following digital biodiversity platforms do you use?   ) *and* The answer was 'I use it in a professional context (before going into the field, in the field during inspections, during data collection and processing, etc.) ' to question '25 [Q17]' (In which context(s) do you use the Observatoire des espèces exotiques envahissantes ? )

***Tick the answer(s) that apply :***

- Required by your management
- Recommended by colleagues
- Recommended by friends/family members
- On your own initiative
- Other:

##### On average, how often do you use the Observatoire des espèces exotiques envahissantes in a professional context ?

Answer this question only if the following conditions are met:

The answer was 'Yes' to question '1 [Q1]' (Have you worked, or do you currently work, in a professional activity related to trees (in towns, green spaces, forests, etc.)? ) *and* The answer was 'Observatoire des espèces exotiques envahissantes' to question '14 [Q14]' (Which of the following digital biodiversity platforms do you use?   ) *and* The answer was 'I use it in a professional context (before going into the field, in the field during inspections, during data collection and processing, etc.) ' to question '25 [Q17]' (In which context(s) do you use the Observatoire des espèces exotiques envahissantes ? )

***Please select one answer below :***

- Every day
- 2 or 3 times a week
- Once a week
- Once a month
- Less than once a month
- Other

##### What type of use do you make of the Observatoire des espèces exotiques envahissantes platform  ?

Answer this question only if the following conditions are met:

The answer was 'Yes' to question '1 [Q1]' (Have you worked, or do you currently work, in a professional activity related to trees (in towns, green spaces, forests, etc.)? ) *and* The answer was 'Observatoire des espèces exotiques envahissantes' to question '14 [Q14]' (Which of the following digital biodiversity platforms do you use?   )

***Tick the answer(s) that apply :***

- I contribute to the platform with my own observations (sharing data on the platform)
- I consult the data (following observations or reports, collecting data)

##### For what purpose do you use the Observatoire des espèces exotiques envahissantes ?

Answer this question only if the following conditions are met:

The answer was 'Yes' to question '1 [Q1]' (Have you worked, or do you currently work, in a professional activity related to trees (in towns, green spaces, forests, etc.)? ) *and* The answer was 'Observatoire des espèces exotiques envahissantes' to question '14 [Q14]' (Which of the following digital biodiversity platforms do you use?   )

***Tick the answer(s) that apply :***

- Identifying species
- Reporting and sharing observations with other users
- Keeping a record of my observations
- Monitoring the presence of pests in a given area
- Other:

##### In which context(s) do you use iNaturalist ?

Answer this question only if the following conditions are met:

The answer was 'Yes' to question '1 [Q1]' (Have you worked, or do you currently work, in a professional activity related to trees (in towns, green spaces, forests, etc.)? ) *and* The answer was ' iNaturalist' to question '14 [Q14]' (Which of the following digital biodiversity platforms do you use?   )

***Tick the answer(s) that apply :***

- I use it in a personal capacity (walks in nature, taking part in a community project, taking part in a family or friends project, etc.)
- I use it in a professional context (before going into the field, in the field during inspections, during data collection and processing, etc.)

##### Is your use of iNaturalist in a professional context :

Answer this question only if the following conditions are met:

The answer was 'Yes' to question '1 [Q1]' (Have you worked, or do you currently work, in a professional activity related to trees (in towns, green spaces, forests, etc.)? ) *and* The answer was ' iNaturalist' to question '14 [Q14]' (Which of the following digital biodiversity platforms do you use?   ) *and* The answer was 'I use it in a professional context (before going into the field, in the field during inspections, during data collection and processing, etc.) ' to question '30 [Q18]' (In which context(s) do you use iNaturalist ? )

***Tick the answer(s) that apply :***

- Required by your management
- Recommended by colleagues
- Recommended by friends/family members
- On your own initiative
- Other:

##### On average, how often do you use iNaturalist in a professional context ?

Answer this question only if the following conditions are met:

The answer was 'Yes' to question '1 [Q1]' (Have you worked, or do you currently work, in a professional activity related to trees (in towns, green spaces, forests, etc.)? ) *and* The answer was ' iNaturalist' to question '14 [Q14]' (Which of the following digital biodiversity platforms do you use?   ) *and* The answer was 'I use it in a professional context (before going into the field, in the field during inspections, during data collection and processing, etc.) ' to question '30 [Q18]' (In which context(s) do you use iNaturalist ? )

***Please select one answer below :***

- Every day
- 2 or 3 times a week
- Once a week
- Once a month
- Less than once a month
- Other

##### What type of use do you make of the iNaturalist platform  ?

Answer this question only if the following conditions are met:

The answer was 'Yes' to question '1 [Q1]' (Have you worked, or do you currently work, in a professional activity related to trees (in towns, green spaces, forests, etc.)? ) *and* The answer was ' iNaturalist' to question '14 [Q14]' (Which of the following digital biodiversity platforms do you use?   )

***Tick the answer(s) that apply :***

- I contribute to the platform with my own observations (sharing data on the platform)
- I consult the data (following observations or reports, collecting data)

##### For what purpose do you use iNaturalist?

Answer this question only if the following conditions are met:

The answer was 'Yes' to question '1 [Q1]' (Have you worked, or do you currently work, in a professional activity related to trees (in towns, green spaces, forests, etc.)? ) *and* The answer was ' iNaturalist' to question '14 [Q14]' (Which of the following digital biodiversity platforms do you use?   )

***Tick the answer(s) that apply :***

- Identifying species
- Reporting and sharing observations with other users
- Keeping a record of my observations
- Monitoring the presence of pests in a given area
- Other:

##### In which context(s) do you use Vigicultures ?

Answer this question only if the following conditions are met:

The answer was 'Yes' to question '1 [Q1]' (Have you worked, or do you currently work, in a professional activity related to trees (in towns, green spaces, forests, etc.)? ) *and* The answer was 'Vigicultures' to question '14 [Q14]' (Which of the following digital biodiversity platforms do you use?   )

***Tick the answer(s) that apply :***

- I use it in a personal capacity (walks in nature, taking part in a community project, taking part in a family or friends project, etc.)
- I use it in a professional context (before going into the field, in the field during inspections, during data collection and processing, etc.)

##### Is your use of Vigicultures in a professional context :

Answer this question only if the following conditions are met:

The answer was 'Yes' to question '1 [Q1]' (Have you worked, or do you currently work, in a professional activity related to trees (in towns, green spaces, forests, etc.)? ) *and* The answer was 'Vigicultures' to question '14 [Q14]' (Which of the following digital biodiversity platforms do you use?   ) *and* The answer was 'I use it in a professional context (before going into the field, in the field during inspections, during data collection and processing, etc.) ' to question '35 [Q19]' (In which context(s) do you use Vigicultures ? )

***Tick the answer(s) that apply :***

- Required by your management
- Recommended by colleagues
- Recommended by friends/family members
- On your own initiative
- Other:

##### On average, how often do you use Vigicultures in a professional context ?

Answer this question only if the following conditions are met:

The answer was 'Yes' to question '1 [Q1]' (Have you worked, or do you currently work, in a professional activity related to trees (in towns, green spaces, forests, etc.)? ) *and* The answer was 'Vigicultures' to question '14 [Q14]' (Which of the following digital biodiversity platforms do you use?   ) *and* The answer was 'I use it in a professional context (before going into the field, in the field during inspections, during data collection and processing, etc.) ' to question '35 [Q19]' (In which context(s) do you use Vigicultures ? )

***Please select one answer below :***

- Every day
- 2 or 3 times a week
- Once a week
- Once a month
- Less than once a month
- Other

##### What type of use do you make of the Vigicultures platform  ?

Answer this question only if the following conditions are met:

The answer was 'Yes' to question '1 [Q1]' (Have you worked, or do you currently work, in a professional activity related to trees (in towns, green spaces, forests, etc.)? ) *and* The answer was 'Vigicultures' to question '14 [Q14]' (Which of the following digital biodiversity platforms do you use?   )

***Tick the answer(s) that apply :***

- I contribute to the platform with my own observations (sharing data on the platform)
- I consult the data (following observations or reports, collecting data)

##### For what purpose do you use Vigicultures?

Answer this question only if the following conditions are met:

The answer was 'Yes' to question '1 [Q1]' (Have you worked, or do you currently work, in a professional activity related to trees (in towns, green spaces, forests, etc.)? ) *and* The answer was 'Vigicultures' to question '14 [Q14]' (Which of the following digital biodiversity platforms do you use?   )

***Tick the answer(s) that apply :***

- Identifying species
- Reporting and sharing observations with other users
- Keeping a record of my observations
- Monitoring the presence of pests in a given area
- Other:

##### If you use other platform(s) not listed here, please indicate them:

Answer this question only if the following conditions are met:

The answer was 'Yes' to question '1 [Q1]' (Have you worked, or do you currently work, in a professional activity related to trees (in towns, green spaces, forests, etc.)? ) *and* The answer was 'I use other platform(s) not mentioned in this list' to question '14 [Q14]' (Which of the following digital biodiversity platforms do you use?   )

***Please write your answer here:………………………………………………***

##### In which context(s) do you use this/these other platform(s) ?

Answer this question only if the following conditions are met:

The answer was 'Yes' to question '1 [Q1]' (Have you worked, or do you currently work, in a professional activity related to trees (in towns, green spaces, forests, etc.)? ) *and* The answer was 'I use other platform(s) not mentioned in this list' to question '14 [Q14]' (Which of the following digital biodiversity platforms do you use?   )

***Tick the answer(s) that apply :***

- I use it in a personal capacity (walks in nature, taking part in a community project, taking part in a family or friends project, etc.)
- I use it in a professional context (before going into the field, in the field during inspections, during data collection and processing, etc.)

##### Is your use of this/these other platform(s)  in a professional context :

Answer this question only if the following conditions are met:

The answer was 'Yes' to question '1 [Q1]' (Have you worked, or do you currently work, in a professional activity related to trees (in towns, green spaces, forests, etc.)? ) *and* The answer was 'I use other platform(s) not mentioned in this list' to question '14 [Q14]' (Which of the following digital biodiversity platforms do you use?   ) *and* The answer was 'I use it in a professional context (before going into the field, in the field during inspections, during data collection and processing, etc.) ' to question '41 [Q20s1]' (In which context(s) do you use this/these other platform(s) ? )

***Tick the answer(s) that apply :***

- Required by your management
- Recommended by colleagues
- Recommended by friends/family members
- On your own initiative
- Other:

##### On average, how often do you use this/these other platform(s)  in a professional context ?

Answer this question only if the following conditions are met:

The answer was 'Yes' to question '1 [Q1]' (Have you worked, or do you currently work, in a professional activity related to trees (in towns, green spaces, forests, etc.)? ) *and* The answer was 'I use other platform(s) not mentioned in this list' to question '14 [Q14]' (Which of the following digital biodiversity platforms do you use?   ) *and* The answer was 'I use it in a professional context (before going into the field, in the field during inspections, during data collection and processing, etc.) ' to question '41 [Q20s1]' (In which context(s) do you use this/these other platform(s) ? )

***Please select one answer below :***

- Every day
- 2 or 3 times a week
- Once a week
- Once a month
- Less than once a month
- Other

##### What type of use do you make of this/these other platform(s)   ?

Answer this question only if the following conditions are met:

The answer was 'Yes' to question '1 [Q1]' (Have you worked, or do you currently work, in a professional activity related to trees (in towns, green spaces, forests, etc.)? ) *and* The answer was 'I use other platform(s) not mentioned in this list' to question '14 [Q14]' (Which of the following digital biodiversity platforms do you use?   )

***Tick the answer(s) that apply :***

- I contribute to the platform with my own observations (sharing data on the platform)
- I consult the data (following observations or reports, collecting data)

##### For what purpose do you use this/these other platform(s)  ?

Answer this question only if the following conditions are met:

The answer was 'Yes' to question '1 [Q1]' (Have you worked, or do you currently work, in a professional activity related to trees (in towns, green spaces, forests, etc.)? ) *and* The answer was 'I use other platform(s) not mentioned in this list' to question '14 [Q14]' (Which of the following digital biodiversity platforms do you use?   )

***Tick the answer(s) that apply :***

- Identifying species
- Reporting and sharing observations with other users
- Keeping a record of my observations
- Monitoring the presence of pests in a given area
- Other:

#### CITIZEN CONTRIBUTIONS TO BIODIVERSITY PLATFORMS

##### In your view, for non-professionals, recognising these tree pests and/or diseases would be:

Answer this question only if the following conditions are met:

The answer was 'Yes' to question '1 [Q1]' (Have you worked, or do you currently work, in a professional activity related to trees (in towns, green spaces, forests, etc.)? ) *and* The answer was 'Yes' to question '10 [Q10]' (Do you take part in the Surveillance of Regulated and Emerging Pests (SORE)? )

|  | Very difficult | Fairly difficult | Fairly easy | Very easy |
| --- | --- | --- | --- | --- |
| Pine wood nematode *(Bursaphelenchus xylophilus)* |  |  |  |  |
| Emerald ash borer *(Agrilus planipennis)* |  |  |  |  |
| Pitch canker *(Fusarium circinatum)* |  |  |  |  |
| Chestnut blight *(Cryphonectria parasitica)* |  |  |  |  |
| Oak processionary moth *(Thaumetopoea processionea)* |  |  |  |  |
| European spruce bark beetle *(Ips typographus)* |  |  |  |  |
| Bronze birch borer *(Agrilus anxius)* |  |  |  |  |
| Pine processionary moth *(Thaumetopoea pityocampa)* |  |  |  |  |
| Canker stain of plane *(Ceratocystis platani)* |  |  |  |  |
| Asian longhorned beetle *(Anoplophora glabripennis)* |  |  |  |  |

##### In your view, the reporting by a non-professional of these tree pests and/or diseases on biodiversity platforms would be :

Answer this question only if the following conditions are met:

(([Q1.NAOK](https://sondages.inrae.fr/index.php/admin/questions/sa/view/surveyid/793573/gid/45011/qid/702614) == "Y") and ([Q10.NAOK](https://sondages.inrae.fr/index.php/admin/questions/sa/view/surveyid/793573/gid/45010/qid/702622) == "Y"))

|  | Very unlikely | Fairly unlikely | Fairly likely | Very likely |
| --- | --- | --- | --- | --- |
| Pine wood nematode *(Bursaphelenchus xylophilus)* |  |  |  |  |
| Emerald ash borer *(Agrilus planipennis)* |  |  |  |  |
| Pitch canker *(Fusarium circinatum)* |  |  |  |  |
| Chestnut blight *(Cryphonectria parasitica)* |  |  |  |  |
| Oak processionary moth *(Thaumetopoea processionea)* |  |  |  |  |
| European spruce bark beetle *(Ips typographus)* |  |  |  |  |
| Bronze birch borer *(Agrilus anxius)* |  |  |  |  |
| Pine processionary moth *(Thaumetopoea pityocampa)* |  |  |  |  |
| Canker stain of plane *(Ceratocystis platani)* |  |  |  |  |
| Asian longhorned beetle *(Anoplophora glabripennis)* |  |  |  |  |

##### How likely are you to consult a biodiversity platform to monitor the presence of these tree pests or diseases?

Answer this question only if the following conditions are met:

(([Q1.NAOK](https://sondages.inrae.fr/index.php/admin/questions/sa/view/surveyid/793573/gid/45011/qid/702614) == "Y") and ([Q10.NAOK](https://sondages.inrae.fr/index.php/admin/questions/sa/view/surveyid/793573/gid/45010/qid/702622) == "Y"))

|  | Very unlikely | Fairly unlikely | Fairly likely | Very likely |
| --- | --- | --- | --- | --- |
| Pine wood nematode *(Bursaphelenchus xylophilus)* |  |  |  |  |
| Emerald ash borer *(Agrilus planipennis)* |  |  |  |  |
| Pitch canker *(Fusarium circinatum)* |  |  |  |  |
| Chestnut blight *(Cryphonectria parasitica)* |  |  |  |  |
| Oak processionary moth *(Thaumetopoea processionea)* |  |  |  |  |
| European spruce bark beetle *(Ips typographus)* |  |  |  |  |
| Bronze birch borer *(Agrilus anxius)* |  |  |  |  |
| Pine processionary moth *(Thaumetopoea pityocampa)* |  |  |  |  |
| Canker stain of plane *(Ceratocystis platani)* |  |  |  |  |
| Asian longhorned beetle *(Anoplophora glabripennis)* |  |  |  |  |

#### TRUST IN DATA FROM DIGITAL BIODIVERSITY PLATFORMS

Please imagine yourself in the following two scenarios:

**SCENARIO 1: While preparing for an inspection, out of curiosity you consult a digital biodiversity platform to find out which species are already present or reported in your area. Browsing the observations on the platform, you spot a photograph showing** [***Anoplophora glabripennis***](https://i0.wp.com/entomologytoday.org/wp-content/uploads/2025/07/asian-longhorned-beetle.jpg?resize=1024%2C983&ssl=1)***,* even though you have never observed this pest in that area.**

***Please indicate how far you agree with the statements below:***

Answer this question only if the following conditions are met:

The answer was 'Yes' to question '1 [Q1]' (Have you worked, or do you currently work, in a professional activity related to trees (in towns, green spaces, forests, etc.)? ) *and* The answer was 'Yes' to question '10 [Q10]' (Do you take part in the Surveillance of Regulated and Emerging Pests (SORE)? )

|  | Strongly disagree | Somewhat disagree | Somewhat agree | Strongly agree |
| --- | --- | --- | --- | --- |
| This information on the presence of Anoplophora glabripennis in your area is reliable |  |  |  |  |
| The absence of observations (photographs, geolocation, taxonomy) of Anoplophora glabripennis on the digital biodiversity platform in other areas means it is genuinely absent there |  |  |  |  |
| The data from this observation (photographs, geolocation, taxonomy) are reliable enough to be used in a professional context |  |  |  |  |
| Digital biodiversity platforms can be useful for early detection |  |  |  |  |
| The number of observations of Anoplophora glabripennis on the digital biodiversity platform is representative of its abundance in the area |  |  |  |  |

**SCENARIO 2: While preparing for an inspection, out of curiosity you consult a digital biodiversity platform to find out which species are already present or reported in your area. Browsing the observations on the platform, you spot a photograph showing** [***Corythuca arcuata***](https://i0.wp.com/insektarium.net/wp-content/uploads/2022/12/C.arcuata-1.jpg?resize=380%2C380&ssl=1) ***,* even though you have never observed this pest in that area.**

***Please indicate how far you agree with the statements below:***

Answer this question only if the following conditions are met:

(([Q1.NAOK](https://sondages.inrae.fr/index.php/admin/questions/sa/view/surveyid/793573/gid/45011/qid/702614) == "Y") and ([Q10.NAOK](https://sondages.inrae.fr/index.php/admin/questions/sa/view/surveyid/793573/gid/45010/qid/702622) == "Y"))

|  | Strongly disagree | Somewhat disagree | Somewhat agree | Strongly agree |
| --- | --- | --- | --- | --- |
| This information on the presence of Corythuca arcuata in your area is reliable |  |  |  |  |
| The absence of observations (photographs, geolocation, taxonomy) of Corythuca arcuata on the digital biodiversity platform in other areas means it is genuinely absent there |  |  |  |  |
| The data from this observation (photographs, geolocation, taxonomy) are reliable enough to be used in a professional context |  |  |  |  |
| Digital biodiversity platforms can be useful for early detection |  |  |  |  |
| The number of observations of Corythuca arcuata on the digital biodiversity platform is representative of its abundance in the area |  |  |  |  |

**How far do you agree with the statement below:**

**Biodiversity platforms make it possible to identify unknown organisms observed in the field effectively, thanks to automatic recognition tools.**

Answer this question only if the following conditions are met:

The answer was 'Yes' to question '1 [Q1]' (Have you worked, or do you currently work, in a professional activity related to trees (in towns, green spaces, forests, etc.)? ) *and* The answer was 'Yes' to question '10 [Q10]' (Do you take part in the Surveillance of Regulated and Emerging Pests (SORE)? )

***Please select one answer below :***

- Strongly disagree
- Somewhat disagree
- Somewhat agree
- Strongly agree

#### Other

##### If you would like to receive the results of our survey, please give your e-mail address:

Answer this question only if the following conditions are met:

The answer was 'Yes' to question '1 [Q1]' (Have you worked, or do you currently work, in a professional activity related to trees (in towns, green spaces, forests, etc.)? )

***Please write your answer here:…………………………………………………………………..***

##### If you would like to add any comments or further details, please write them here:

Answer this question only if the following conditions are met:

The answer was 'Yes' to question '1 [Q1]' (Have you worked, or do you currently work, in a professional activity related to trees (in towns, green spaces, forests, etc.)? )

***Please write your answer here:…………………………………………………………………***

**Thank you for the time you have given to our survey. Your participation helps to advance research on the role of citizen science in tree and forest health surveillance.**

**Thank you for completing this questionnaire.**
